# Genetic and pharmacologic alterations of claudin9 levels suffice to induce functional and mature inner hair cells

**DOI:** 10.1101/2023.10.08.561387

**Authors:** Yingying Chen, Jeong Han Lee, Jin Li, Seojin Park, Maria C. Perez Flores, Braulio Peguero, Jennifer Kersigo, Mincheol Kang, Jinsil Choi, Lauren Levine, Michael Anne Gratton, Bernd Fritzsch, Ebenezer N. Yamoah

## Abstract

Hearing loss is the most common form of sensory deficit. It occurs predominantly due to hair cell (HC) loss. Mammalian HCs are terminally differentiated by birth, making HC loss challenging to replace. Here, we show the pharmacogenetic downregulation of *Cldn9*, a tight junction protein, generates robust supernumerary inner HCs (IHCs) in mice. The ectopic IHC shared functional and synaptic features akin to typical IHCs and were surprisingly and remarkably preserved for at least fifteen months >50% of the mouse’s life cycle. *In vivo*, *Cldn9* knockdown using shRNA on postnatal days (P) P2-7 yielded analogous functional ectopic IHCs that were equally durably conserved. The findings suggest that Cldn9 levels coordinate embryonic and postnatal HC differentiation, making it a viable target for altering IHC development pre- and post-terminal differentiation.

## Introduction

Mammalian cochlear hair cells (HCs) comprise a single row of inner hair cells (IHCs) and three rows of outer hair cells (OHCs). HCs transduce sound-mediated mechanical force into neural electrical codes for ear-brain intercommunication. The IHCs are the predominant afferent transducers, while the OHCs amplify low-level sound. Mammalian HCs are terminally differentiated by birth, and they are susceptible to damage by ototoxic drugs, noise-overexposure, aging, and environmental insults, resulting in hearing loss, the most common sensory deficit (Neitzel and Fligor, 2019; Rybak and Ramkumar, 2007; Wilson and Tucci, 2021; Wu et al., 2020). Emerging understanding of the mechanisms of transcription factors that induce HC differentiation (Atkinson et al., 2014; Bermingham et al., 1999; Garcia-Anoveros et al., 2022; Zheng and Gao, 2000), and potential induction for regeneration are promising, but none have produced new HCs with sustained functions (Iyer et al., 2022b; Zine et al., 2021). Thus, the current treatment for hearing loss ensuing from HC loss is cochlear implants, despite the potential advantages of HC-replacement therapy (Hinton et al., 2021; Sullivan et al., 2020).

In the mammalian cochlea, each HC is separated from the next by intervening supporting cells (SCs), forming an invariant and alternating mosaic along the cochlea’s length. Cochlear SCs can divide and trans-differentiate into HCs in the neonatal stage, serving as a potential resource for HC differentiation using transcription and developmental signaling factors (White et al., 2006). At the neonatal stage, *Atoh1*, a basic helix-loop-helix factor, induces SCs trans-differentiation into HCs. Upregulation of *Atoh1, GFI1,* and *POU4F3* triggers HC differentiation, but the fledgling HCs invariably degenerate, suggesting pre-maturity (Iyer et al., 2022a; Liu et al., 2012). Additionally, inhibition of the Notch signaling or upregulation of the wnt1 pathway suffices to drive HC formation from SCs (Tang et al., 2023), but the functional features of the newly developed HCs are circumspect (Mizutari et al., 2013; Waqas et al., 2016). While transcription factors are potent targets for developmental regulation, a cadre of these essential DNA-binding proteins and the precise timing of expression are required to complete the developmental cascade (Jahan et al., 2015). Thus, the key is identifying HC-developing bands of transcription and signaling factors to treat hearing loss. Recent studies identified the transcription factors INSM1 and IKZF2 as the regulators of OHC fate, while the transcription factor TBX2 specifies and maintains HC and SC fate (Kaiser et al., 2022; Li et al., 2023), advancing understanding of HC-subtype developmental specification mechanisms.

Contact-mediated lateral inhibition is among the final developmental events, where once a cell fate is determined, it inhibits neighboring cells from becoming that cell type. The HC-SC interphase is laced with tight junction proteins (TJPs), which may mediate lateral inhibition mechanisms in nonmammalian vertebrates, though their additional function is unclear in mammals (Chrysostomou et al., 2012; Goodyear and Richardson, 1997; Petrovic et al., 2014). In addition, TJPs were also found to regulate cell proliferation (Bhat et al., 2020; Díaz-Coránguez et al., 2019). The TJP scaffold cingulin regulates lateral inhibition in HC-SC rearrangements in the avian basilar papilla (Goodyear and Richardson, 1997), and claudin b (*cldnb*), an ortholog of the *cldn4* in humans, upregulation controls cellular patterning during HC regeneration in zebrafish (Montalbano et al., 2021). Moreover, damaged HC extrusion and the breaking of intercellular junctional adhesions may trigger differentiation and regenerative proliferation (Corwin et al., 2007). In mammals, E-cadherin’s junctional expression negatively alters HC regenerative capacity (Burns et al., 2008; Burns et al., 2013), while *Cldn9* is a positive regulator of cell proliferation (Hong et al., 2014; Zavala-Zendejas et al., 2011). Although the high expression of *Cldn9* in the organ of Corti (OC) is well-established (Kudo et al., 2018; Nakano et al., 2009) (**Fig. 1A-C**), it is unknown whether regulation of the TJP contributes to sensory cell differentiation.

**Figure 1.**
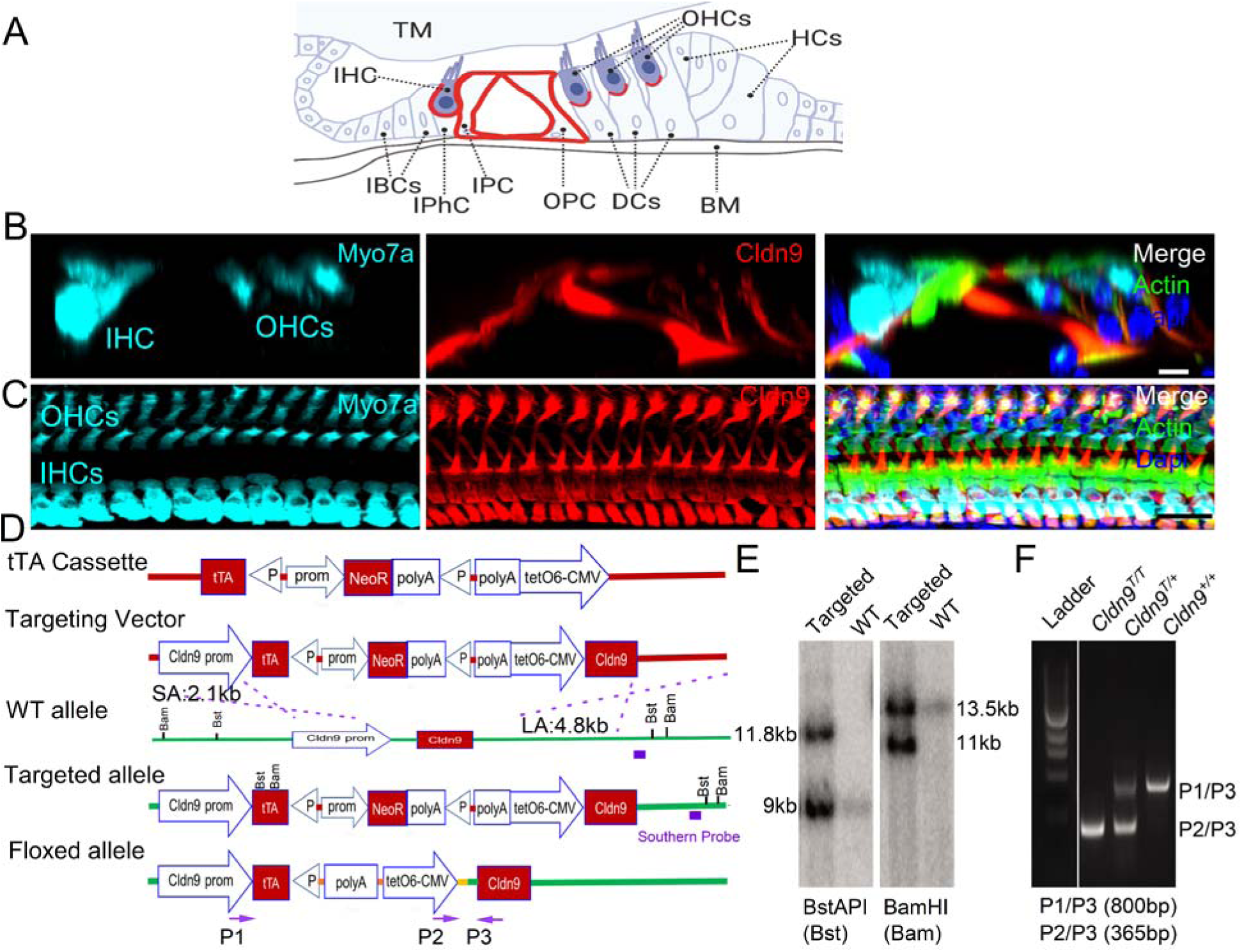
The cochlear partition and construct of doxycycline-tet-OFF-*Cldn9* mouse line. **A.** Schematic diagram of the radial section of the cochlear partition. The organ of Corti (OC) is seated on the basilar membrane (BM). The inner and outer pillar cells (IPC and OPC) are separated from the inner and outer hair cells (IHC, OHCs) by tight junction proteins (TJPs). The red lines indicate the expression outline of Cldn9. TM: tectorial membrane; IHC: inner hair cell; OHCs: outer hair cells; IBC: inner border cell; IPhC: inner phalangeal cell; IPC: inner pillar cell; DCs: Deiters cells; HCs: Hensen’s cells. **B.** The immunostaining of *Cldn9* in the 8-wk old WT (*Cldn9^+/+^*) mouse cochlea showing IHC stained with myosin7a (cyan) and claudin9 (red)antibodies, phalloidin, actin (green) for stereocilia bundles and Dapi (blue) for nuclear stain. **C**. Surface view of the cochlea **B**. Scale = 10 *μ*m. **D.** The construct of Doxycycline-tet-OFF- *Cldn9* in the ES cells. Generation of a genetically modified mouse line in which the level of *Cldn9* gene expression can be regulated by doxycycline: A tetracycline-based genetic switch (tTA cassette) consists of three main modules, the tetracycline-controlled transcriptional activator (tTA), the neomycin resistance gene flanked by LoxP sites, and the final module that contains six copies of the tet operator (tetO) fused to the minimal CMV promoter. The tTA cassette was inserted at the −110-nucleotide position upstream of the translational start of *Cldn9* to generate the targeting vector. The targeting vector was electroporated into B6 mouse embryonic stem cells. Following the selection in G418, DNA samples from the neomycin-resistant ES cell clones were prepared for short-arm PCR/sequencing analysis and Southern blot analysis to confirm the insertion of the tTA cassette in the ES cells. The genetically modified ES cells containing one copy of the tTA cassette were injected into the healthy albino B6 blastocysts, and the injected blastocysts were transplanted into the uterus of an albino B6 mouse to generate the chimeric mouse. The chimeric mouse was then bred with albino B6 mice to produce the F1 heterozygous mouse, and the germline transmission was confirmed by tail DNA genotyping. The deletion of the selection marker in the tTA cassette by crossing the F1 mouse with the embryonic Cre line (B6.129S4-Meox2 tm1(cre)Sor /J). **E**. Genomic DNA samples were prepared from neomycin-resistant ES cell clones and digested with BstAPI or BamHI. The tTA cassette insertion was confirmed by the Southern blot analysis. **F**. Mouse tail DNA genotyping was carried out using primers P1/P3 for WT and P2/P3 for targeted alleles showing *Cldn9^T/T^*, *Cldn9^T/+^* and *Cldn9^+/+^*samples.

To determine the roles in vivo of Cldn9, we generated doxycycline (dox)-tet-OFF-Cldn9 transgenic mice to regulate expression levels of Cldn9. The downregulation of Cldn9 resulted in functional supernumerary (ectopic) putative IHCs along the cochlear contour. Auditory neurons innervated ectopic mechanically transducing IHCs with synaptic features resembling normal IHCs. Analogous additional putative IHCs differentiation was observed when Cldn9-shRNA was injected through the round window to postnatal (P) days (P2, P4, and P7) mice, suggesting that regulation of Cldn9 levels coordinates embryonic and postnatal development differentiation of SC into IHCs. Notably, the ectopic IHCs at the apical and middle-frequency contour of the cochlea were preserved for over half the mouse’s life cycle (15 months), making Cldn9 a viable target for generating supernumerary IHCs.

## Results

To control *Cldn9* levels *in vivo*, we generated a mouse model with a site-specific genetic switch regulated by dietary doxycycline (dox) and dox-containing drinking water without interfering with the typical profile of *Cldn9* expression ((Supplement figure 1 (**S1**)). **Figures 1D-F** show the design constructs and Southern blot analysis to confirm the insertion cassette in the ES cells. The genotyping was performed using PCR with the tail tissue. Transgene was generated on mixed C57/B6 and backcrossed into a CBA-CaJ (CBA) background after 12 generations to reduce accelerated progressive hearing loss (Peguero and Tempel, 2015; Sergeyenko et al., 2013; Willott and Erway, 1998). Results of quantitative RT-PCR from the three groups of animals, including wild-type littermates (*Cldn9^+/+^)*, heterozygote *(Cldn9^+/T^),* and homozygous (*Cldn9^T/T^)* with (1 mg/ml) and without dox treatment. Without dox treatment, *Cldn9^T/T^* and *Cldn9^+/T^*mice demonstrate ∼55 and a 40-fold increase in *Cldn9* mRNA expression in cochlear tissue compared with *Cldn9^+/+^* cochleae (**S2**). Treatment of *Cldn9^+/T^* mice with dox (1 mg/ml) resulted in an ∼0.4-0.6-fold decline in mRNA levels compared to *Cldn9^+/+^* cochleae (**S2**), translating to a marked difference in Cldn9 protein expression (**S2**). Immunoelectron microscopic analysis showed that Cldn9 levels reduced by ∼8-fold in the *Cldn9^+/T^* cochlea (**S3**). The *Cldn9^T/T^* mice had a reduced survival rate (1.5±0.3 months) relative to the *Cldn9^+/T^* littermates (23±2 months). Thus, all experiments were restricted to *Cldn9^+/T^ and Cldn9^+/+^* littermates fed on dox-water (1 mg/ml). There were no recognizable differences in body weight between *Cldn9^+/T^* and *Cldn9^+/+^* mice for both females and males (**S2**). All animals were in the CBA background. *Cldn9* downregulation in the *Cldn9^+/T^* cochlea showed a qualitative decrease in the Cldn6 and an increase in ILDR1 TJP levels but no comparative differences in others (Kitajiri et al., 2004; Kitajiri and Katsuno, 2016) (**S4**).

### Downregulation of *Cldn9* induces the production of ectopic cochlear HCs

5-week-old mice *Cldn9^+/T^* cochleae displayed a notable row of ectopic HCs (**Fig. 2A-E)**. The ectopic HCs were observed along the cochlear contour, ranged in abundance from base to apex (**S5**), and had contact with innervating neurons, shown in cochlear sections (**Fig. 2A)**. The ectopic HCs were positively labeled with anti-myosin VIIa antibody and phalloidin-labeled stereocilia bundles, which are features of typical HCs. Moreover, the ectopic HCs were not labeled with antibodies to prestin (**S6**), a marker for OHCs, suggesting the new HCs were likely derived from the IHC lineage. A distinctive "U"-shaped IHC bundle was apparent (**Fig. 2D)**. Scanning electron microscopic (SEM) was used for high-resolution analyses to evaluate IHC and their bundle morphology. The original and ectopic IHCs in *Cldn9^+/T^*mice had normal morphology, including intact hair bundles (**Fig. 2E**). However, the stereocilia bundle orientation was less orderly when compared to those in *Cldn9^+/+^* cochlea (**Fig. 2D-E**). In addition to their shape, the ectopic HCs expressed multiple CtBP2 labeled synaptic structures in contrast to typical OHCs (**Figs. 3B, 5B**), and reacted positively to otoferlin antibodies (Pangrsic et al., 2010; Roux et al., 2006; Strenzke et al., 2016) (data not shown). We denote the new HCs as "ectopic" IHCs. IHC counts at different ages (P2-P21) and the cochlear frequency segments (8 kHz and 32 kHz) demonstrate that the *Cldn9*-induced ectopic IHCs were most prominent at the cochlear apex but remained statistically significant at the base (**Figs 2E-F**). Viable ectopic IHCs were identified in 15-month *Cldn9^+/T^* mice in the apical cochlea (**Fig. 3**). For OHCs, the numbers along the cochlear contour, apex, middle, and base were not significantly different among the two genotypes at P14 (**S7A**) and at the apical turn of organ of Corti in 15-month-old *Cldn9^+/T^* mice (**S7B**).

**Figure 2.**
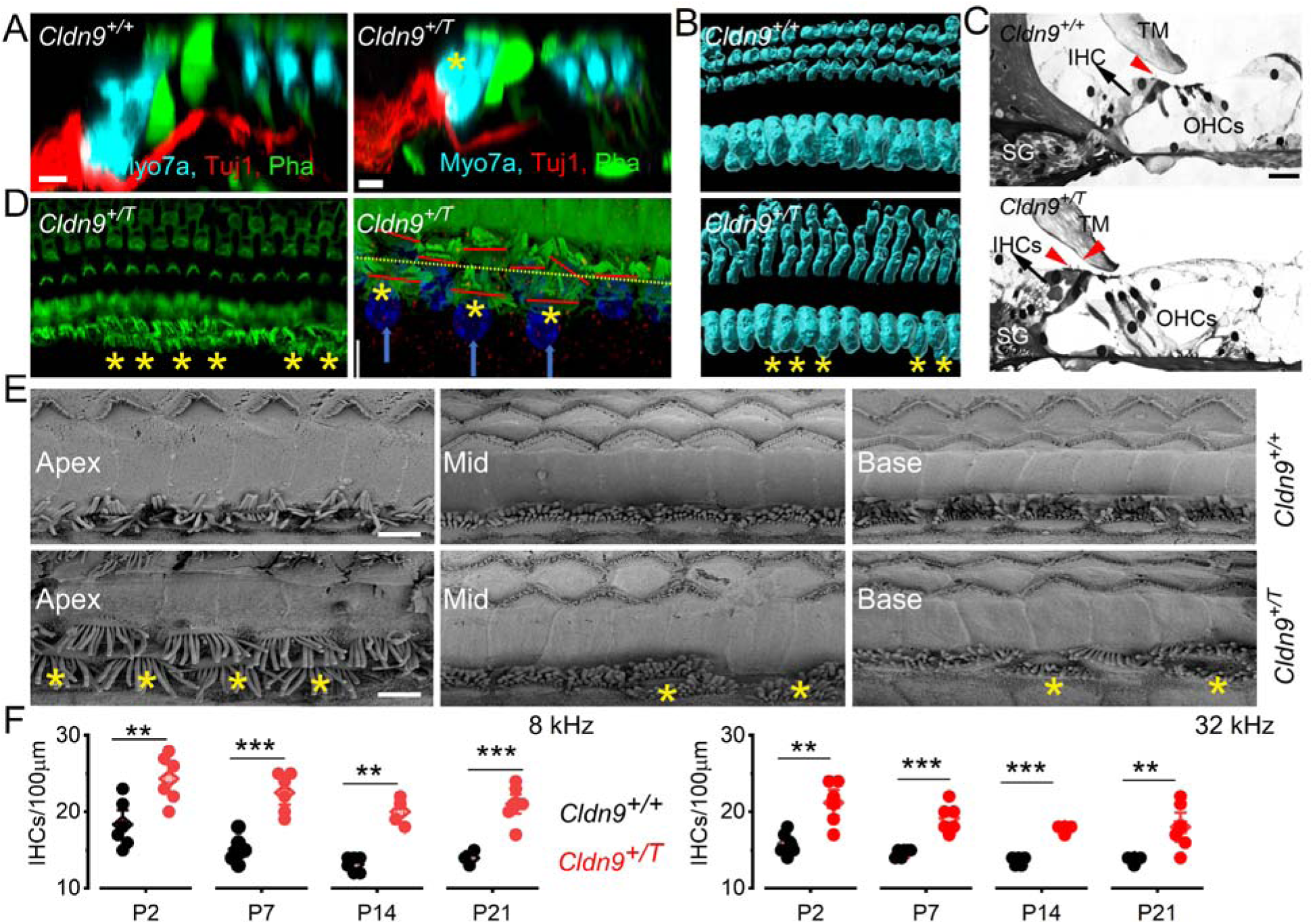
*Cldn9* knockdown-induced supernumerary cochlear IHCs. **A**. Radial section of the cochlea partition of 8-wk-old WT (*Cldn9^+/+^*) (left Panel) and *Cldn9*+/T (right Panel) mice. Myosin 7a (cyan) stained IHCs and phalloidin (green) labeled actin stereocilia bundles and other cellular structures, and Tuj1 (red) stained neurons. The yellow asterisks (*) denote ectopic-IHC. Scale bar = 5 *μ*m. **B.** 3-D rendition of cochlear surface preparation (aged 8-wks old), showing a single row of IHCs and three rows of OHCs in the WT (top Panel: *Cldn9^+/+^*) and for the *Cldn9^+/T^*(lower Panel). The ectopic IHCs are marked with (*) Scale bar = 10 *μ*m. **C.** Mid-modiolar section of the 8-week-old cochlea of *Cldn9^+/+^*(top) and *Cldn9^+/T^* (lower) panels. The red arrows show one and two IHC bundles in the *Cldn9^+/+^* and *Cldn9^+/T^*cochlear sections, respectively. SG = spiral ganglion, TM= tympanic membrane IHC = inner hair cell, OHCs = outer hair cells Scale bar = 10 *μ*m. **D.** Surface preparation of cochlea of *Cldn9^+/T^*, showing two-rows of IHCs (the 2^nd^-row IHCs are marked with (*) and higher magnification of a tilted section revealing the nuclei (Dapi stain in blue, and arrow points to the nuclei of the 2^nd^ IHC row). The dotted yellow line shows the hair bundles of the 1^st^ row, and the red lines denote the orientations of the hair bundles. Scale bar = 10 *μ*m. **E.** Scanning electron photomicrographs of the apical (Apex), middle (MT), and basal (Base) cochlear turn of 6-week-old *Cldn9^+/+^*and *Cldn9^+/T^* mice. The ectopic IHCs are marked with (*). **F.** Summary data of Quantification of IHCs count at 8- and 32-kHz placed cochlear map at postnatal days (P) 2, 7, 14, 21 in *Cldn9^+/+^*compared with *Cldn9^+/T^* mice. Each data point is the means of 4-blind counts. At 8 kHz cochlear placed-map, the *p* values were P2 = 7.4X10^-3^, P7 =3.0X10^-4^, P14 = 2X10^-3^, and P21 = 1.5X10^-4^. At 32 kHz cochlear placed-map, the *p* values were P2 = 4.0X10^-3^, P7 =1.0X10^-3^, P14 = 8.5X10^-6^, and P21 = 1.7X10^-2^.

**Figure 3.**
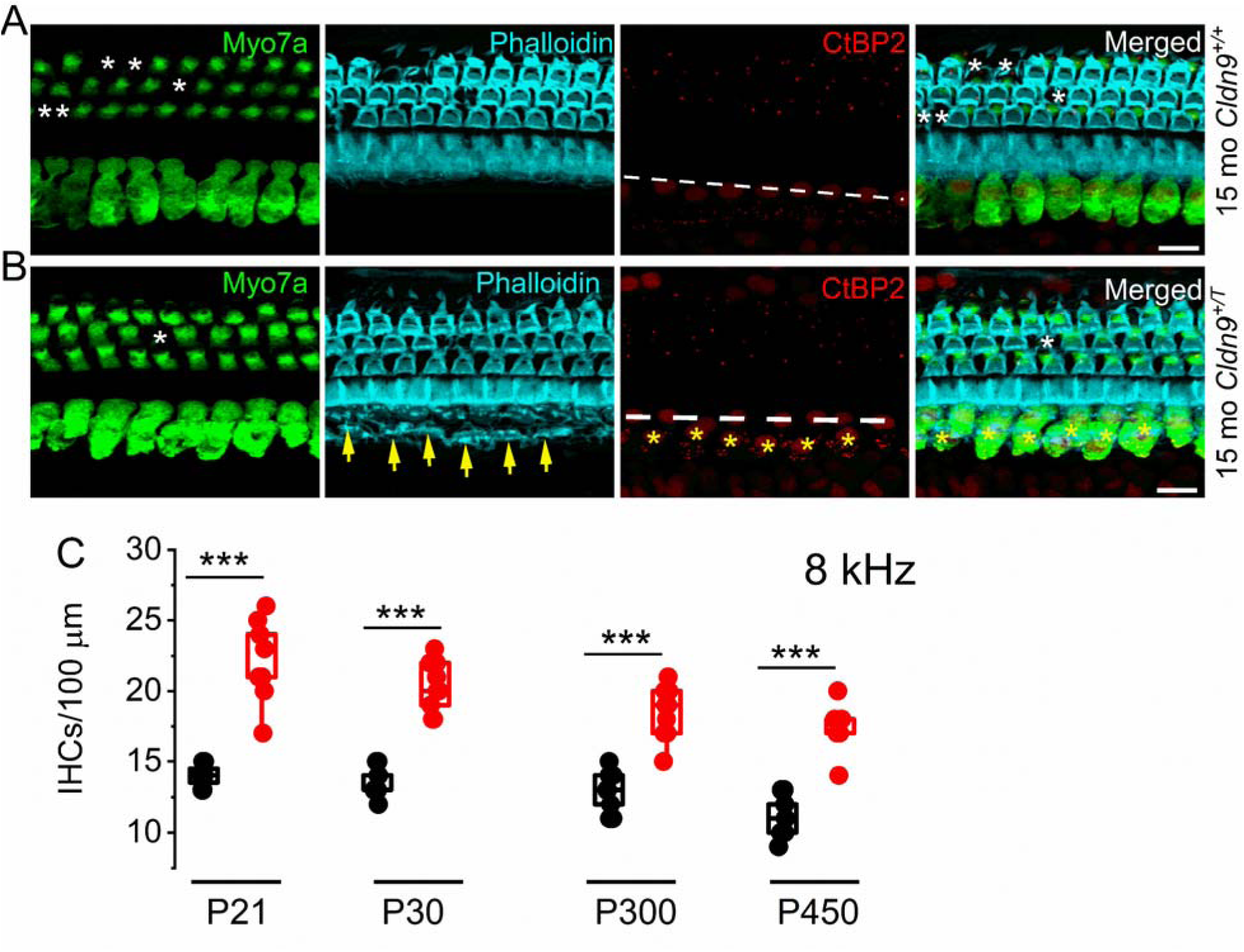
*Cldn9* knockdown-induced supernumerary IHCs remain intact in older mice. **A.** Surface cochlear preparation of a 15-months (P450) old *Cldn9^+/+^* mouse shows IHCs and OHCs labeled with Myosin7a (Myo7a) antibody (green) and actin with phalloidin (cyan) and presynaptic maker CtBP2 antibody (red), which is nuclear-positive. The white dash line marks the single row of IHCs. Note that the missing OHCs are marked with white asterisks at 15 months. **B.** Similar preparation as in **A**, from a 15-month-old *Clnd9^+/T^* mouse. Viable ectopic IHC can be seen with yellow arrows pointing to the hair bundles and nuclei, stained with CtBP2 antibody (*). **C.** Summary data of Quantification of IHCs count at 8- and 32-kHz placed cochlear map at postnatal days (P) 21, 30, 300, and 450 in *Cldn9^+/+^* compared with *Cldn9^+/T^*mice. Each data point is the mean of 3-blind counts. At 8 kHz cochlear placed-map, the *p* values were P30 = 2.9X10^-7^, P300 =3.7X10^-6^, and P450 = 1.4X10^-7^. At 32 kHz cochlear placed-map, the *p* values were P30 = 1.1X10^-2^, P300 =4.6X10^-4^, and P450 = 6.7X10^-5^.

### Functional features of *Cldn9*-induced ectopic IHCs resemble normal IHCs

Motor responses of both mouse genotypes’ (*Cldn9^+/^*^+^ and *Cldn9^+/^*^T^) to auditory stimuli (Preyer’s test) were normal. To evaluate the status of IHC function, we analyzed auditory brainstem responses (ABR) to various sound-pressure levels. *Cldn9^+/T^*mice responses exhibited similar characteristic responses to broadband clicks and pure tones at 8, 16, and 32 kHz stimuli (**Fig. 4A**), with ∼5-15 dB threshold elevation in the *Cldn9^+/T^* mice [*F* (12, 579) =2.07, *P*=0.02]. The pattern of hearing threshold remained virtually constant from 2-8 months of monitoring [*F* (12, 319) =0.50, *P*=0.91]. Moreover, *Cldn9^+/^*^+^ and *Cldn9^+/T^*mice yielded similar distortion products [*F* (6, 357) =0.45, P=0.84] (**Fig. 4B**), suggesting normal OHC function. Besides, a significant drop of ABR wave 1 amplitude was observed in the 2 months *Cldn9^+/^*^T^ mice comparing to the wild-type littermates (8 kHz *F* (10, 235)=2.91, *P*=0.02; 16 kHz *F* (10, 237)=1.99, *P*=0.04; 32 kHz *F* (10, 233)=7.80, P=0.00; click *F* (10, 233)=3.99, *P*=0.00 (**S8A-D**). However, wave 1-latency shows no difference between *Cldn9^+/^*^T^ and *Cldn9^+/^*^+^ mice at 2 months (**S8E-H**). The apparent normal morphology of the ectopic IHCs led to the hypothesis that the IHCs may exhibit functional mechano-electrical transducer (MET) currents. Since active MET channels are partially open at rest, the rapid uptake of the channel permeable lipophilic dye FM1-43 (Gale et al., 2001), was assessed. Results were determined from FM1-43 dye loading of apically located original and ectopic IHCs in 4-week-old *Cldn9^+/T^* cochlear site representing ∼4-6-kHz characteristic frequencies (CFs). Local perfusion of 10-*μ*M FM1-43 resulted in intense dye labeling at the hair bundle level of original and ectopic IHCs. The dye-membrane partitioning and diffusion across the IHCs’ basal aspects occurred within seconds. Z-stacked-time-lapse images were taken below hair bundle level 2-sec post dye exposure and at the basolateral compartment (**Fig. 4C**). The time constants (*τ*) of dye loading at the bundle and supra-nuclear basal membrane levels for original IHCs were (data from 4 mice); 19.4±1.1 sec (n=27) and 29.0±3.8 sec (n=27), and for ectopic IHCs were 23.1±4.5 sec (n=27) and 47.0±8.4 sec (n=27) (**Fig. 4D**). These results are consistent with functioning IHCs at rest. We conclude that the ectopic IHC bundles are set at the optimum dynamic range to transduce MET current at rest like the original counterparts. IHC MET current magnitudes and kinetic profiles from the original and ectopic rows at the ∼3-4-kHz cochlear-place map in P21 mice were comparable, as summarized in figures **4E-G**. The normalized current-displacement relationships were well-fitted with a two-state Boltzmann function, portraying marked similarities between the original and the ectopic IHC MET currents (**Fig. 4H**). However, the displacement-response relationship for the ectopic IHCs was right-shifted, indicative of reduced sensitivity.

**Figure 4.**
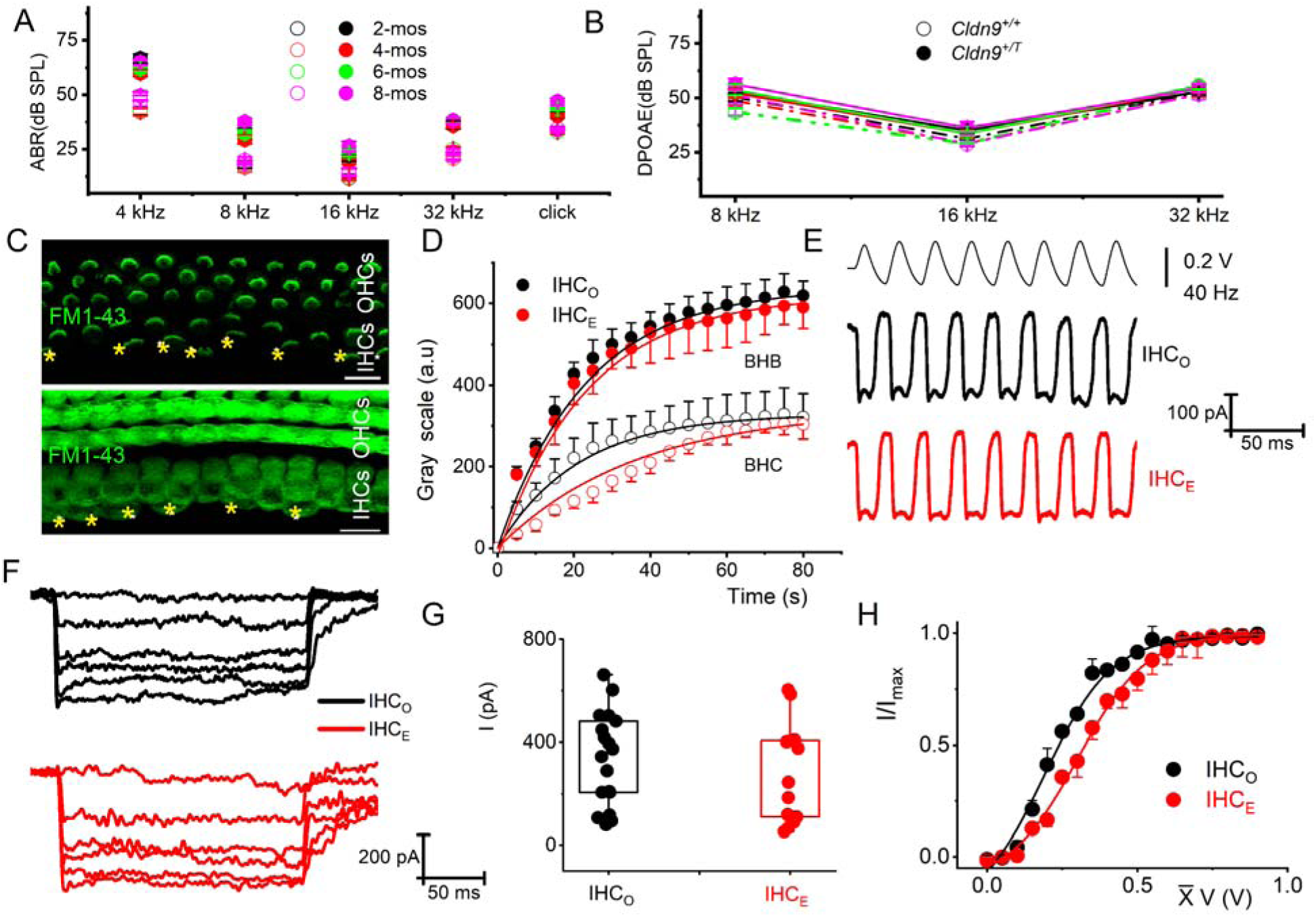
Auditory brainstem recordings (ABR) and distortion product otoacoustic emissions (DPOAE) assessment and mechanoelectrical transducer channel-FM1-43 permeation and currents elicited from original (O; IHC_O_) and ectopic (E; IHC_E_) IHC stereocilia bundles. **A,** ABR thresholds from the *Cldn9^+/+^* (open symbols, das lines) and *Cldn9^+/T^* (solid symbols, solid lines) monitored from 2-8 months. The sound pressure levels (SPL) in dB of pip tones (in kHz) 4, 8, 16, and 32 and broadband clicks delivered to the ear are indicated. ABR thresholds for *Cldn9^+/+^* (n = 12) and *Cldn9^+/T^* (n = 17) mice. On average, the *Cldn9^+/T^* had a 5-15 dB elevated threshold compared to *Cldn9^+/+^* mice. Within each group of mice, there was no significant shift in ABR threshold from 2-8 months of monitoring. The data means ± SD. **B,** Mean DPOAE threshold for 2-8 months (*Cldn9^+/+^* group: n=9; *Cldn9^+/T^* group: n = 14) was tested, measuring the 2f1 - f2 DPOAE over a geometric-mean frequency range at 8, 16, and 32 kHz. The lines and symbols are the same as the labels in Fig. 4A. On average, the *Cldn9^+/T^* had a 5-7 dB elevated threshold compared to *Cldn9^+/+^* mice. **C,** Fluorescent images of FM1-43 were taken at different time points at two focal planes: the base of IHC bundle (BHB) and the below cuticular plate (BHC) levels of postnatal day (P) 14 *Cldn9^+/T^*cochlea. The ectopic IHC bundles are marked with (*). **D.** Time 0 indicates the onset of dye application (10 mM for the 5-sec duration). Consecutive images at BHB and BHC were taken at 5-s intervals. Dye loaded into this region remained sustained. The dye enters the apical aspects of the cell before being visualized at the basal pole. (Scale bar, 10 *μ*m). The change in fluorescence at focal levels as a function of time (adjusted for the interval between frame capture at each level). Densitometric data of mean pixel intensity were measured in arbitrary grayscale units. Summary data from IHC_O_ and IHC_E_ (n = 27 from 3 mice). Frames were taken from 3-4 kHz placed map of the mouse cochlea. The change in fluorescence was fitted with an exponential function, and the time constants (t, in s) of FM1-43 dye loading in IHC_O_ (in black circles) at BHB and BHC levels,19.4±1.1 (n = 27) and 29.0±3.8 (n = 27). Similar analyses for IHC_E_ (in red circles) at BHB and BHC levels were 23.1±4.5 (n = 27) and 47.0±8.4 (n = 27). **E,** Typical MET current traces from IHC_O_ and IHC_E_ elicited sinusoidal hair bundle deflection at 40 Hz. **F.** MET current elicited a series of ∼200-ms hair bundle displacements, using fluid-jet deflection towards the taller stereocilia. Hair cells were held at −80 mV. All recordings were made from apical IHCs. Bundle deflection was elicited with 0.1-0.9 V pressure clamps in 0.1-V steps. Estimates for the exact bundle displacement were not determined. For clarity, a few traces were omitted. **G,** Summary of group data of the maximum IHC MET current measured from IHC_O_, 335±181 pA (n = 18) and IHC_E_, 268±192 (n = 14) (mean ± SD) **H,** Normalized displacement response relationships fitted with a two-state Boltzmann function. The half-maximum displacement voltage (in V) for IHC_O_ and IHC_E_ were 0.20±0.02 and 0.30±0.01 (n= 7).

### Synaptic features of ectopic IHCs and original IHCs

To determine whether the ectopic IHCs had additional properties in terms of systemic functions, we examined features such as neuronal innervation in both the original and ectopic IHCs. We labeled auditory neurons and IHCs with calretinin (Calb2) antibodies (Sun et al., 2018). Results show Calb2-positive-subtype neurites at the modiolar aspects of both IHCs (**Fig. 5A**). The synapses between the IHCs and auditory neurons at the apical, middle, and basal cochlear locations from 5-week-old *Cldn9^+/+^* and *Cldn9^+/T^* mice show substantial differences. The organization of afferent synapses was significantly different, identified as paired presynaptic-CtBP2 (red) and postsynaptic-Homer1 (green) immunopuncta. Contrasting *Cldn9^+/+^* from *Cldn9^+/T^* cochlear samples, results showed reduced synaptic numbers in the *Cldn9^+/T^* (**Fig. 5B-C**). Moreover, quantifying the mean number of synapses per IHC among the original and ectopic IHCs showed variations along the cochlear axis (**Fig. 5D**). This data suggests that ectopic IHCs are equipped to serve as afferent receptors capable of transducing and transmitting mechanical displacement into neural codes, while synaptic transmission from ectopic IHCs might be compromised because significant differences in synapse numbers were identified.

**Figure 5.**
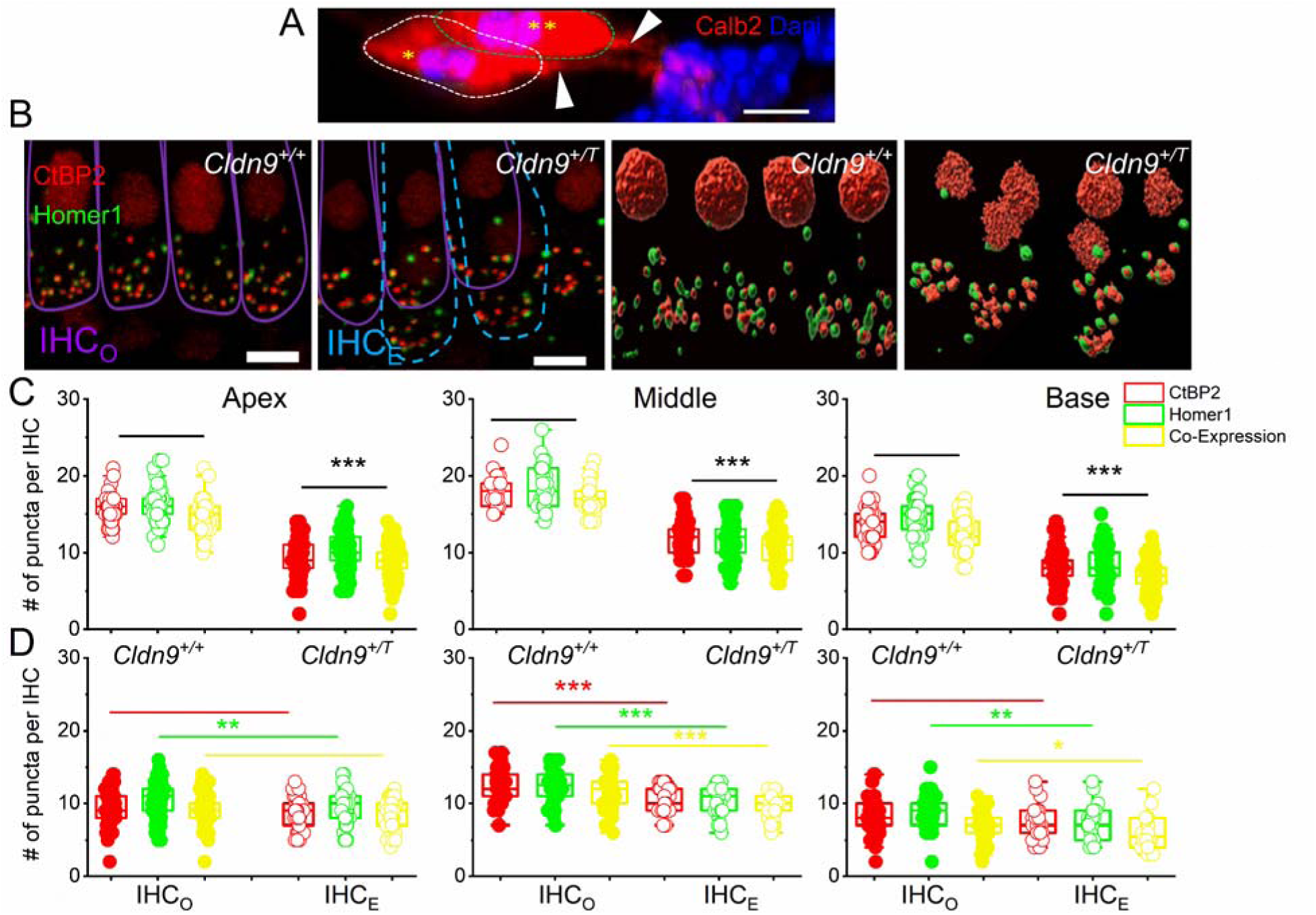
Pre- and post-synaptic features of original and ectopic IHCs in *Cldn9^+/+^* and *Cldn9^+/T^* mice. **A,** The innervation of calretinin (Calb2)-positive afferent neurons (red) of an original and ectopic IHCs (marked *, and **) from a 5-wks-old *Cldn9^+/T^*mouse cochlea. Dapi (blue) labeled the nuclei. Arrows point to the nerve terminals. Scale bar = 5 *μ*m. **B,** CtBP2 labeled (red) pre-synapse and Homer1 labeled (green) post-synapse in the cochlear IHCs in the *Cldn9^+/+^* and *Cldn9^+/T^* mice. The original IHC (IHC_O_) and ectopic IHC (IHC_E_) outlines are marked with purple and cyan dashed lines. The two right panels show a 3-D rendition of the pre-and post-synaptic markers in *Cldn9^+/+^* and *Cldn9^+/T^* cochleae. Scale bar = 5 *μ*m. **C,** Quantification of the number of pre- (red) and post-synaptic (green) immunopuncta at three tonotopic cochlear locations: apex (3-5 kHz), middle (12-16 kHz), and base (32-40 kHz) from 8-wk-old *Cldn9^+/+^*(open circles) and *Cldn9^+/T^* (solid circles) mice. Colocalized pre- and post-synaptic marker counts per IHC are shown in yellow. Values (mean ± SD) are illustrated (*p* < 0.05, 0.01, 0.001 = *, **, ***). Comparing pre-, post-synaptic and colocalized markers for apical IHCs, between *Cldn9^+/+^* (n = 56 from 3 mice) and *Cldn9^+/T^* (n = 117 from 4 mice) the *p* values were pre- *p* = 2.2X10^-42^, post-=3.5X10^-30^, co-expression, 4.3X10^-34^. For mid-cochlea IHCs, between *Cldn9^+/+^*(n = 29 from 3 mice) and *Cldn9^+/T^* (n = 79 from 4 mice) the *p* values were pre- *p* = 1.7X10^-19^, post- =1.3X10^-14^, co-expression, 2.5X10^-18^. For basal-cochlea IHCs, between *Cldn9^+/+^*(n = 97 from 3 mice) and *Cldn9^+/T^* (n = 94 from 4 mice) the *p* values were pre- *p* = 3.5X10^-27^, post- =8.2X10^-27^, co-expression, 2.9X10^-26^. **D,** analysis of synaptic features among IHC_O_ and IHC_E_ in *Cldn9^+/T^* cochlea. At the cochlea apex, comparing pre- post- and colocalized markers between IHC_O_ (n = 76 from 3 mice) and IHC_E_ (n = 41 from 3 mice), the *p* values were pre- *p* = 7.0X10^-2^, post- = 9.0X10^-3^, co-expression = 7.0X10^-2^. For mid-cochlea comparing pre-, post- and colocalized markers between IHC_O_ (n = 48 from 3 mice) and IHC_E_ (n = 30 from 3 mice) the *p* values were pre- *p* = 1.4X10^-4^, post- = 8.9X10^-5^, co-expression, 1.7X10^-5^. For basal-cochlea comparing pre-, post- and colocalized markers between IHC_O_ (n = 65 from 3 mice) and IHC_E_ (n =22 from 3 mice) the *p* values were pre- *p* = 9.0X10^-2^, post- = 1.0X10^-2^, co-expression, 5.0X10^-2^.

### Postnatal induction of ectopic IHCs by shRNA knockdown

Because pregnant mothers were fed on dox-water from gestation, the ectopic IHCs in *Cldn9^+/T^* cochlea were of embryonic origin. We designed an efficient shRNA construct to knock down *Cldn9* postnatally. Viral transduction *in vivo*, through round window injection into the cochlear tissue, was monitored with a GFP reporter gene (**Fig. 6A**). We first assessed the efficiency and specificity of the shRNA knockdown of *Cldn9* using quantitative RT-PCR (**S9A**) and immunofluorescence microscopy (**Fig. 6A**). Four days post-injection, there was an ∼20-fold reduction in *Cldn9* mRNA relative to nontargeting scrambled shRNA injected cochlea (**Fig. 6A, S9**). Transduction *in vivo* of *Cldn9* shRNA into P2-7 inner ears yielded cochleae with ectopic IHCs compared to internal controls, consisting of opposite cochlea injected with nontargeting scrambled shRNA injected, which did not display ectopic IHCs. By contrast, the P14 inner ear injected with *Cldn9*-shRNA produced no detectable increase in ectopic IHCs as counted by three independent blinded observers (**Fig. 6B-D**). Ultrastructural SEM analysis of *Cldn9*-shRNA transduced P2-7 inner ears show ectopic IHC with hair bundles resembling original IHCs (**Fig. 6C**). The MET currents invoked from the ectopic-IHCs induced by postnatal *Cldn9* knockdown were in keeping with functional HCs.

**Figure 6.**
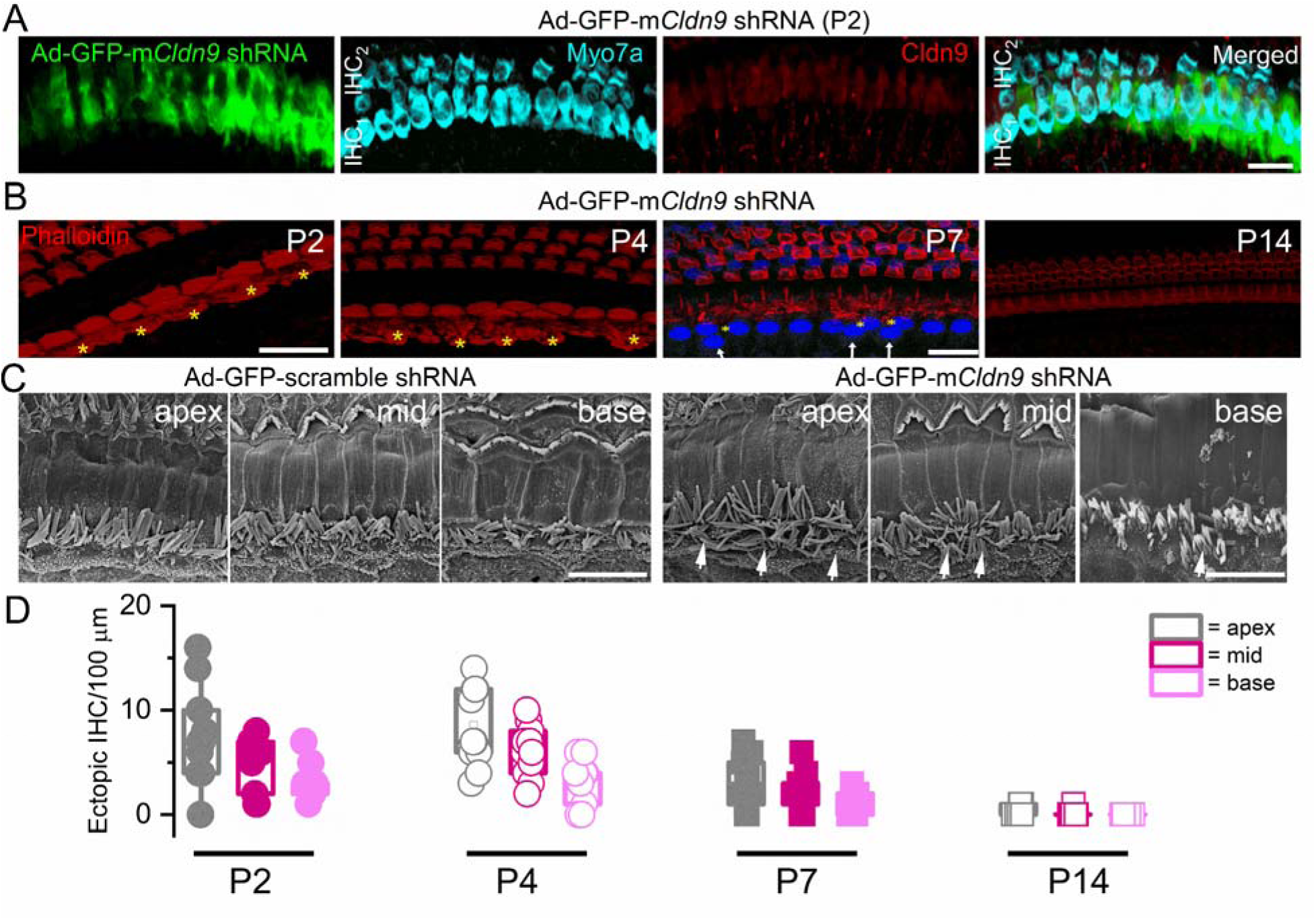
Postnatal induction of ectopic IHCs with the Cldn9 *shRNA* knockdown. **A**, Ectopic IHCs observed in P21 cochlea after *Cldn9 shRNA* injection at P2. Downregulation of *Cldn9* levels by ∼20-fold compared to scrambled shRNA is shown in S7A, and a reduced protein expression (Fig. 6A) is shown in the 3^rd^ panel (*Cldn9*, red). GFP (green) expression is a marker for adeno (AD)-virus-transduced cells. The right panel shows the merged image with two rows of IHCs stained with HC marker Myo7a (cyan) and Ad-GFP-mCldn9 shRNA (green) expression in the adjacent row of supporting cells. The Ad-GFP-mCldn9 shRNA (green) is expressed in HCs. **B,** Examples of phalloidin stained whole-mount organ of Corti samples from cochleae of the wild-type mice injected at P2, P4, P7, and P14 with Cldn9 shRNA. The tissues were harvested 20 days after viral injection. Sections at or near the cuticular-plate level show actin labeled with phalloidin (red). Several ectopic IHCs are marked with asterisks in yellow (*). Where nuclei were labeled with Dapi (blue), we marked the ectopic IHC nuclei with a white arrow. **C,** Scanning electron photomicrographs of the apical (apex), middle (mid), and basal (base) cochlear turn of 5-6-wks-old mice after scrambled shRNA and *Cldn9* shRNA injection at P2, showing original and ectopic IHCs (marked with white arrows). Images were captured at the cochlea’s apical, middle, and basal contour. **D,** Summary data of quantification of ectopic IHCs count at cochlear apex, middle, and base for samples where *Cldn9* shRNA was injected at P2, P4, P7, and P14. Each data point (n=11) is the mean of 3-blind counts from 3 mice. Ectopic IHCs at-P2 injection (mean/100-mm±SD) for apical, middle, and basal cochlea were 7.5±4.5, 5.0±2.4, and 3.2±1.6; at-P4; 8.5±3.8, 6.1±2.5, and 3.2±1.6; at-P7; 2.8±2.4, 2.4±1.7, and 1.1±1.0; at-P14, 0.2±0.5, 0.1±0.3, and 0.0±0.0. Data are plotted to show individual replicates (n=11; meanlJ±lJSD). There were significant differences at *p*<0.05 level for comparison between P2-7 and P14 F (11,120) = (17) *p* = *4.2X10^-20^*. *Post hoc* comparisons using the Tukey HSD test indicate that post-shRNA injection-induced ectopic IHCs are significantly different at P2 apex vs P14 apex (*p*=2.4X10^-8^); P2 mid vs. P14 mid (*p*=1.3X10^-4^); and P2 base vs. P14 base (*p*=7.0X10^-2^).

### The endocochlear potential and K^+^ concentrations in *Cldn9^+/T^* mice

The cochlear duct is furnished with cellular syncytia, K^+^ channels, and transporters/pumps that operate to orchestrate a unidirectional (basal to apical) flux of K^+^ at the lateral wall to produce the endocochlear potential (EP, ∼+80 mV), an extracellular potential, subserving the proverbial powerhouse for HC functions (Von Bekesy, 1952). A remnant of K^+^ flux is a high K^+^ endolymph (∼140 mM) restricted from leakage into the basolateral aspects of HCs by TJ between HCs and SCs. At four months of age, the magnitude of the EP and the K^+^ concentration of the endolymph and perilymph of *Cldn9^+/T^* mice were variable significantly from age-matched littermates. A slight decline in the amplitude of the EP and a substantial rise in perilymph K^+^ was detected in 8-month-old *Cldn9^+/T^,* but there is no statistical difference compared with the detection in 4-month-old Cldn9^+/T^(**S10**). It is unclear whether the modest changes in the EP and K^+^ concentration of perilymph can account for the threshold increase in the *Cldn9^+/T^*mice.

## Discussions

Fate determination is typically completed by birth in cochlear HCs, the primary receptors for mechanosensory sound detection. Generating functional HCs in the mammalian cochlea, with a proper cellular organization that allows for cochlear sound frequency selectivity, has been a demanding yet unsolved challenge. Consequently, deafness resulting from HC loss, which constitutes a significant portion of sensorineural hearing loss, is incurable. Multiple aspects must be considered in inducing HC differentiation since HCs have distinct functional and structural features along the cochlear axis. For optimal sensing of different sound frequencies, cochlear apical-to-basal HCs have diverse structural configurations and ion channel configurations and densities that sculpt and preserve low-to-high frequency sound processing (Kimitsuki et al., 2001). Additionally, HCs are connected to SCs by TJ proteins, enhancing the sensitivity of cochlear OHC sound amplification and maintaining high K^+^ concentration in the endolymph at the apical surface and low K^+^ concentration in the perilymph milieu surrounding HCs (Cohen-Salmon et al., 2002). This process establishes a unidirectional K^+^ flux in the cochlear duct to generate the EP, which boosts the receptor potential of HC by ∼80 mV (Hudspeth, 2008; Von Bekesy, 1952). Thus, besides overcoming the insurmountable terminal differentiation of HCs, newly differentiated HCs *in vivo* should be equipped with features that resemble the original primary HCs to integrate into the specialized cochlear environment.

Among the multiple approaches used with limited success in HC replacement are overexpressing transcription factors involved in prosensory cell differentiation and silencing inhibitory factors in the induction of HC fate. These methods include 1) Transfecting proneural genes, such as *Atoh1*, in embryonic, newborn, and damaged mature cochlear tissue. However, HC-competent cells invariably lose their responsiveness post-birth and in adult animals. In the case of the damaged cochlea, it is challenging to distinguish between transdifferentiated and repaired HCs (Gubbels et al., 2008; Izumikawa et al., 2005; White et al., 2006). 2) Simulating cell division using cell-cycle inhibitors, this strategy activates apoptotic genes, leading to cell death and deafness (Lowenheim et al., 1999). 3) Inhibiting Notch signaling using a *γ*-secretase inhibitor to stimulate HC differentiation from potential inner ear resident stem cells after noise trauma (Jeon et al., 2011; Mizutari et al., 2013). The scarcity of resident stem cells in adult cochlea may limit this strategy. Thus, multiple therapeutic armamentariums are required to restore hearing in the translational setting.

Previous studies demonstrated that claudin-9 is essential for hearing function and the maintenance of auditory HCs, using an ethyl nitrosourea-induced *Cldn9* mutant mouse model (Nakano et al., 2009), which resulted in OHC degeneration. The current results demonstrate that embryonic regulation of *Cldn9* levels, but not null deletion using the dox-tet-OFF-*Cldn9* transgenic strategy induces functional ectopic IHCs, but not OHCs, along the cochlear contour with increasing numbers from base to apex. These *Cldn9* downregulated-induced IHCs mature, acquiring robust MET currents and neural innervation with synaptic structures markedly like resident IHCs. Results also revealed that postnatal downregulation of *Cldn9* levels *in vivo*, using shRNA, suffice to coordinate SC differentiation into IHCs. Because the ectopic IHCs remain viable for a sizable duration of the mouse’s lifespan, the Cldn9 regulatory strategy to induce IHC differentiation subserves a feasible approach to replace lost HCs. The downregulation of Cldn9-mediated selective IHC-increase indicates Cldn9’s role during the latter phase of HC differentiation, perhaps post-OHC-fate determination (Garcia-Anoveros et al., 2022). The findings suggest Cldn9-mediated effects may be upstream of the transcriptional factor-mediated trans-differentiation of HCs since the ectopic HCs had features of IHCs, in contrast to primordial HCs generated by *Atoh1* (Yang et al., 2012). It deepens our understanding of the importance of tapping into later stages of HC differentiation, which will likely result in end-organ-specific HCs.

Moreover, a pragmatic strategy requires titrated levels of the TJP to render new HCs without compromising the sensory epithelial cellular syncytial, a decline in the EP, and, significantly, a gradual extracellular K^+^ increase that mediates undue HC depolarization and death (Nakano et al., 2009). The critical period at which alteration of TJP level can induce ectopic and new IHCs remains unclear, although, in the current report, we have demonstrated that functional and viable mature ectopic IHCs can be generated by regulating Cldn9 levels.

Our findings that downregulation in the TJP, Cldn9, can regulate IHC differentiation are in conceptual agreement with reports demonstrating that lateral inhibition can affect HC specification (Lanford et al., 1999; Stone and Rubel, 1999). Similar accounts are described where newly formed HCs express delta1-like (DII1) and jagged 2 (Jag2) ligands to mediate Notch1-receptor activation in adjacent antecedent cells, thereby inducing the expression of hairy and enhancer of split (Hes1/5), which suppresses pro-HC transcription factors (Bermingham et al., 1999; Chrysostomou et al., 2012). Consistent with this scheme, interruption of Notch1 signaling during HC development leads to HC overproduction (Brooker et al., 2006; Lanford et al., 1999; Zine et al., 2000). In lateral inhibition, the foremost developing cells adopting HC fate antagonize the neighboring cells from differentiating into HCs through direct cell-to-cell communication (Brown and Groves, 2020; Mizutari et al., 2013). A possible explanation for the current findings is that TJ proteins, mainly Cldn9, are signaling in Notch-mediated lateral induction (Daudet and Lewis, 2005; Lewis, 1998). Canonical Notch signaling is activated when a Notch ligand, such as Delta-like1 (DI1) in an adjacent HC, binds to a receptor in the SC, resulting in the release of the intracellular domain of the Notch receptor (NICD), which translocate to the nucleus to activate the Notch-targeted gene transcriptions. The findings also confirm that the Notch signaling pathway is responsible for homeostatic TJP expression *in vitro* and promotes barrier function *in vivo* in the RAG1-adoptive transfer model of colitis (Mathern et al., 2014). Indeed, occluding junction depletion disrupts Notch and mitogen-activated protein kinase (MAPK) signaling in intestinal tissue (Fairchild et al., 2016). In the scheme provided in **Figure 7**, Cldn9 may subserve the signaling catalyst to activate NICD cascades that suppress neighboring SCs from trans-differentiation. A limitation of the model is that if *Cldn9*-induced effects were solely dependent on the Notch signaling, ectopic OHCs and IHCs would have ensued. Alternatively, *Cldn9* levels disruption may alter the mechanical properties of the developing and maturing organ of Corti that may trigger ectopic IHC differentiation, an epiphenomenon independent of the Notch signaling. Future studies and emerging findings on HC differentiation will likely address these shortcomings (Kaiser et al., 2022; Li et al., 2023). In cochlear tissue, downregulation of Cldn9 led to concomitant reduced expression of Cldn6 and increased ILDR1. It is unclear whether the induction of ectopic resulted from reduced expression of *Cldn9* alone or combined TJP alterations. It is conceivable that targeting a different TJP may have similar effects on OHC differentiation, requiring impending studies.

**Figure 7.**
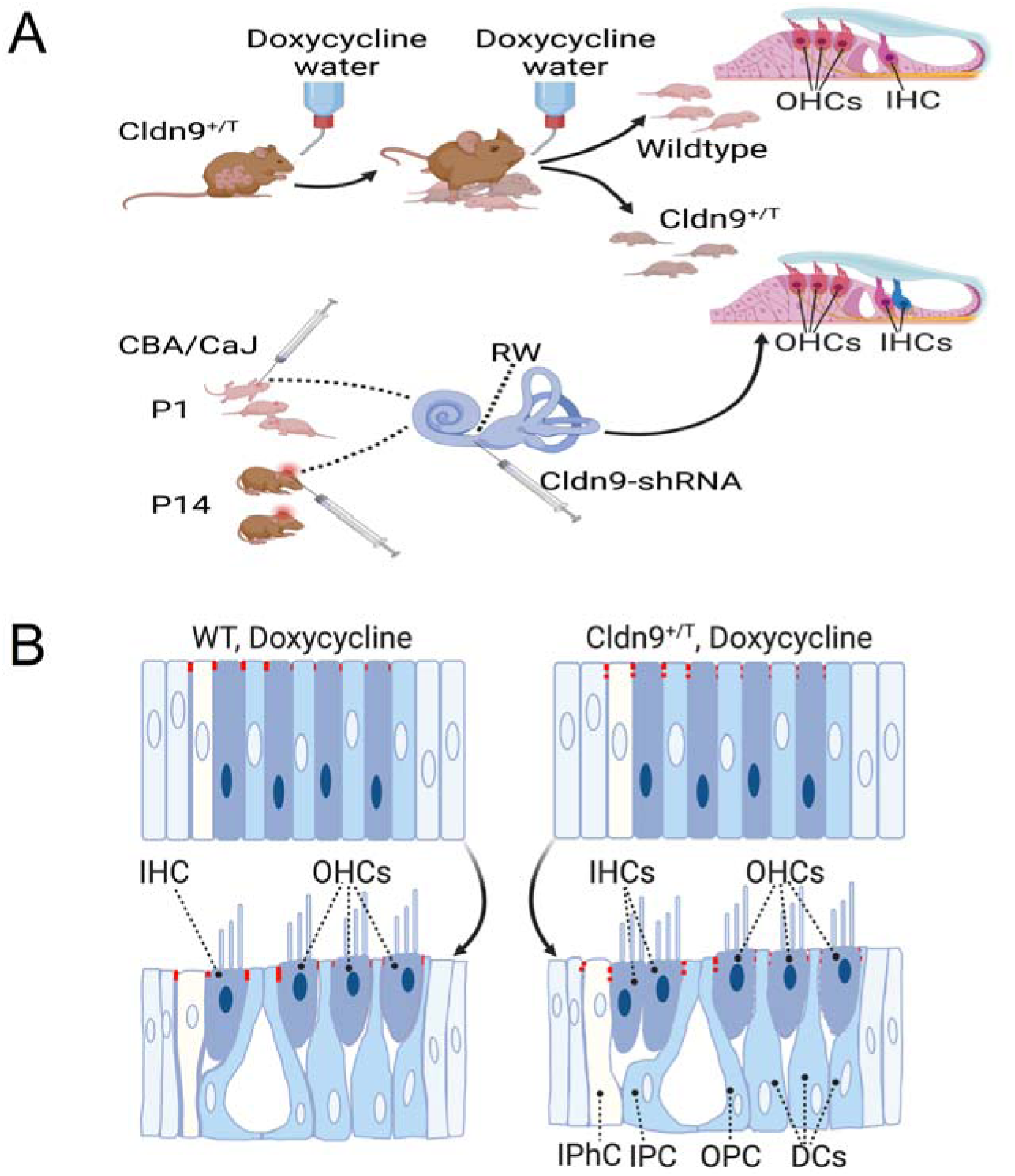
Study design and summary of findings. **A,** The study design schematic included the doxycycline-treated *Cldn9^+/T^* and *Cldn9^+/+^*mice and postnatal mice transduced with *Cldn9* shRNA. Whereas ectopic IHCs were generated by inner ear injection of Cldn9 shRNA from P2-7, we did not detect significant ectopic IHCs at P14. **B,** The contact lateral inhibition in the late-differentiation stage of IHCs and SCs. The solid red line showed the intact Cldn9 sealing between HCs and SCs, while the dashed red line showed the downregulated *Cldn9* level between cells.

## Materials and Methods

All procedures were performed under research guidelines of the institutional animal care and use committee of the University of Nevada, Reno. Mice of either sex were studied. Doxycycline (dox)-tet-OFF-Cldn9 transgenic mice were generated. In the mouse line, dox concentration can regulate the level of Cldn9 gene expression. The construct consisted of a tetracycline-based genetic switch (tTA cassette) made of three main modules: 1) The tetracycline-controlled transcriptional activator (tTA); 2) The neomycin resistance gene flanked by LoxP sites; and 3) Six copies of the tet operator (tetO) fused to the minimal CMV promoter. The tTA cassette was inserted at the −110 -nucleotide position upstream of the translational start of *Cldn9* to generate the targeting vector. The targeting vector was electroporated into B6 mouse embryonic stem cells. Following the selection in G418, DNA samples from the neomycin-resistant ES cell clones were prepared for short-arm PCR/sequencing analysis and Southern blot analysis to confirm the insertion of the tTA cassette into ES cells. Genetically modified ES cells containing one copy of the tTA cassette were injected into healthy albino B6 blastocysts, and the injected blastocysts were transplanted into the uterus of an albino B6 mouse to generate the chimeric mouse. The chimeric mouse was then bred with albino B6 mice to produce the F1 heterozygous mouse, and the germline transmission was confirmed by tail DNA genotyping. The selection marker was deleted in the tTA cassette by crossing the F1 mouse with the embryonic Cre line (B6.129S4-*Meox2^tm1(cre)Sor^*/J). We backcrossed the B6/129S4 background unto the CBA/CaJ mouse background for 12 generations to prevent the masking of age-related hearing loss effects. 1.0 mg/ml of dox water were fed to *Cldn9* breeding pairs from breeding day one, and heterozygote *(Cldn9^+/T^)* and Homozygous (*Cldn9^T/T^)* mice and wild-type littermates (*Cldn9^+/+^)*, through the time until for the sample collections. The body weights of mice were recorded. Genotyping was performed by PCR using a set of primers that flank the knockin in the *Cldn9* gene: forward primer Cldn9 knockin-F (knockin-sequence): 5’–ATCCACGCTGTTTTGACCTC–3’, Cldn9 R3 (Reverse): 5’– TCTGGACCACACAGGACATC– 3’. PCR fragments were separated with 2% agarose gel for an 800 bp product in wild-type, 365 bp in homozygous mutants, and two in heterozygous littermate mice (**Fig. 1**).

### Auditory brainstem recordings (ABR) and Distortion product otoacoustic emissions (DPOAE) measurements

*Cldn9* mice (*Cldn9^+/+^*, *Cldn9^+/T^*, and *Cldn9^T/T^*) littermates were tested at 2-8 months of age. Mice were anesthetized with ketamine and xylazine by intraperitoneal (IP) injection (25 mg/kg). Body temperature was monitored using a rectal probe and maintained at 36.8±1.0°C using a homeothermic device (Harvard Apparatus). ABR and DPOAE measurements were described previously (Dou et al., 2004). For ABR assays, thresholds were obtained by presenting tone bursts at 4, 8, 16, and 32 kHz and a clicking sound from 0 dB to 90 dB sound pressure levels (SPL) in 5 dB intervals. Tones were 2.5 ms, while click was 0.1 ms in duration, with a repetition rate of 21/s. Electrodes were placed sub-dermally behind the tested ear (reference), the vertex (active), and the back (ground). Evoked potentials were averaged over 512 repetitions and collected using a Tucker Davis Technology (TDT) RZ6 processor and BioSigRZ software. The threshold was defined as the lowest intensity of stimulation that yielded a repeatable waveform based on an identifiable ABR wave.

DPOAE measurements were performed using the same TDT system with two calibrated MF1 speakers connected to an ER10B+ microphone. Data was collected every 21 ms and averaged 512 times. DPOAEs were recorded using two pure tones with frequencies f1 and f2, using an f2/f1 ratio of 1.2. Input/output (I/O) functions were obtained by increasing the primary tone L1 (and corresponding L2) in 5-dB steps from 20 to 80 dB SPL at 8, 16, and 32 kHz frequencies. During DPOAE testing, the probe assembly was placed in the mouse’s left ear canal after visual inspection to ensure no ear infection or inflammation of the tympanic membrane. DPOAE thresholds were defined as the lowest level of f1 required to produce a DPOAE ≥ −5 dB SPL(Chen et al., 2021).

### Cochlear mapping and hair cells and synaptic counts

The cochlea was micro-dissected into three to five pieces following the method described (Montgomery and Cox, 2016). Cochlear pieces were measured, and a frequency map was computed based on a 3D reconstruction of the sensory epithelium for HCs and synapse count of associated structures to relevant frequency regions using a custom plug-in to ImageJ (Muniak et al., 2013). Confocal z-stacks of the 4, 8, 16, and 32 kHz areas were collected using a Leica Stellaris8 (Leica) and Nikon A1R laser scanning confocal microscope (Nikon Instruments Inc.). Images were gathered in a 512 x 512 raster using a high-resolution oil immersion objective (60x). IHCs and OHCs at the frequency locations were quantified using myosin-VIIa-positive as an HC-marker within a 70-100-μm field (Chen et al., 2021). Synaptic ribbons and homer-labeled puncta could be counted manually or automatically by setting the filters for the spots function using 3D (x-y-z axis) representations of each confocal z-stack with the microscopic image analysis software Imaris (Oxford Instruments, USA).

### Inner ear histological analysis

The cochleae were intra-labyrinthine perfused through the oval and round windows with 4% paraformaldehyde (PFA). The samples were decalcified in 10% EDTA up to 72-96 hrs, depending on the age, at 4°C. Micro dissected pieces were immunostained with antibodies to the following: (1) mouse anti-C-terminal binding protein 2 (pre-synaptic-marker, BD Biosciences, 1:200, Cat # 612044), (2) rabbit anti-myosin-VIIa (HC-marker, Proteus Biosciences, Inc,1:600, Cat # 25-6790), (3) mouse anti-sox2 (supporting cell (SC)-marker, Santa Cruz Biotechnology, Inc, 1:200, Cat # sc-365823), and (4) rabbit anti-Homer 1 (post-synaptic marker, Synaptic Systems, 1:250, Cat # 160 003), (5) rabbit anti-immunoglobulin like domain containing receptor 1 (ILDR1) (Antibodies-online.com, 1:200, Cat # ABIN1386369), (6) mouse anti-Cldn9 (Santa Cruz, 1:200, Cat # sc-398836), (7) mouse anti-Cldn6 (Santa Cruz, 1:200, Cat # sc-393671), (8) rabbit anti-Sox2 (Abcam, 1:200, Cat # ab97959), (9) rabbit anti-Tuj1 and chicken anti-Tuj1 (Abcam,1:500, Cat # ab18207, ab41489), (10) goat anti-calretinin (Swant Inc., 1:500, code # CG1), (11) mouse anti-calretinin (Millipore Sigma, 1:200, Cat # MAB1568), (12) rabbit anti-Prestin (Santa Cruz, 1:200, Cat # sc-22692), and (13) rabbit anti-calbindin (Cell signaling technology, 1:200, Cat # 13176S) with appropriate secondary antibodies coupled to Alexa-405, −488, −568, and −647 fluorophores.. DAPI labeled the cell nucleus after secondary antibody incubation. Samples were stained with phalloidin and mounted with Fluoro-Gel (Electron Microscopy Sciences). Images were captured with a confocal microscope.

### RNA extraction and quantitative RT-PCR of Cochlear tissue

The cochlea was dissected from the mouse and homogenized on ice. Because of limited tissue, we combined 10-15 mice cochleae for the study. Total RNA was isolated using the RNeasy Plus Mini Kit (Qiagen), and cDNA was generated using the RT2 First Strand Kit (Qiagen). cDNA was combined with RT2 SYBR Green Master Mix (Qiagen), specific qRT-PCR primers, and qRT-PCR analysis was run using the ViiaTM 7 Real-Time PCR System (ABI). Primer efficiencies were determined by standard dilution curve analysis. Three separate samples were used from 10 animals for each group. The experiments from each sample were performed in triplicate, and average cycle threshold (Ct) values were normalized to GAPDH expression. ΔΔCt values were determined relative to *Cldn9^+/+^* cochlear samples. Fold change was defined as 2(−ΔΔCt). Primers used include Gapdh (SA Biosciences) and Cldn9 (ThermoFisher).

### Electron Microscopy

Transmission electron microscopy (TEM) and scanning electron microscopy (SEM) of cochlear sensory epithelia were performed as described (Schwander et al., 2007; Senften et al., 2006). Five to eight-week-old *Cldn9* mice and littermates were sacrificed for TEM, and the cochleae were fixed in 2.5% glutaraldehyde in 0.1 M cacodylate buffer at 4°C overnight. After several washes with buffer alone, cochleae were fixed in 1% osmium tetroxide at RT for 1-hr. After that, the fixed cochleae were decalcified in 10% EDTA for 3–4 days. Fixed and decalcified cochleae were dehydrated using a graded ethanol series and embedded in epoxy resin. Ultrathin sections were cut with a diamond knife. Specimens were examined using an electron microscope.

For SEM, mice were perfused with 4% PFA in 1x PBS, inner ears were isolated, and the stapes footplate was removed. Ears were flushed and fixed overnight in 4% PFA and 2.5% glutaraldehyde in 1x PBS. After washing in ddH_2_O 3X for 1 hour, the samples were post-fixed with 1% osmium tetroxide for approximately 1 hour. Samples were washed before decalcifying for 3-4 days in 0.25 M EDTA at 4°C with daily solution changes. The cochleae were micro dissected, the tectorial membranes removed, and gradually dehydrated in 30%, 50%, 70%, 80%, 90%, 100% ethanol, 2:1 ethanol/hexamethyldisilazane (HMDS, Thermo Scientific #A15139.AE), 1:2 ethanol/HMDS, and finally 100% HMDS. Samples were transferred to an open well plate in HMDS and allowed to air dry overnight in a fume hood. They were then mounted on aluminum stubs (Ted Pella #16111) using double-sided carbon tape (EMS #77817-12) and stored in a specimen mount holder (EMS #76510) sealed in a desiccator until sputter-coated with Au/Pd (Emitech Sputter Coater K550) and viewed. Images were captured utilizing the Hitachi S-4800 SEM. An accelerating voltage of 1kV and 5kV was used. Images were compiled using CorelDRAW X7 graphic suite software.

### Measurements of the endocochlear potential (EP) and K^+^ concentration

*Cldn9* heterozygote mice and wild-type littermates were anesthetized using ketamine and xylazine (100/25 mg/kg, i.p.) and K^+^ concentration, and the EP was measured using double-barreled microelectrodes. An incision was made along the midline of the neck, and soft tissues were bluntly dissected laterally to expose the trachea and the animal’s left bulla. A tracheostomy was done, and the musculature over the bulla was cut posteriorly to expose the bone. A small hole was made in the cochlear capsule directly over the scala media of the lower basal turn. The EP electrode was filled with 300 mM NaCl, the K^+^-selective barrel was silanized, and the tip was filled with a liquid ion exchange (Fluka 60398, K^+^ ionophore I-Cocktail B) that was backfilled with 150 mM KCl. A round-window approach made measurements in the basal turn of the cochlea through the basilar membrane of the first turn. The K^+^-selective electrode was calibrated in solutions with known cation (K^+^ and Na^+^) concentrations *in situ* at 37°C.

### shRNA CLDN9 knockdown

siRNAs were designed, using siRNA at WHITEHEAD software, and cloned under U6 promoter in pSilencer5.1-U6 vector to produce hairpin siRNAs (shRNAs; Vector Biolabs, NM_020293). shRNA was packaged into adeno-associated virus constructs: AAV/Anc80L65-GFP-U6-mCLDN9-shRNA (4) at 1.0X10^12^ GC/mL titer. shRNA sequence for mouse CLDN9 is 5’-CACC GTGCTTCGGGACTGGATAAGACTCGAGTCTTATCCAGTCCCGAAGCAC TTTTT-3’. 1.0 µl shRNA or scrambled shRNA was injected into the round window of mice at P2 – 7 and P14. Mice were anesthetized by induced hypothermia and kept on a cold surface during the injection procedure. After disinfecting the skin with 70% ethanol and Povidone iodine, an incision was made in only the left ear (experiment) and right ear (scramble shRNA injection). Underlying fat and soft tissue were carefully dissected to expose the round window of the cochlea. The glass pipette was pulled with a P-2000 (Sutter Instrument, Novato, CA) and sharpened with a BV-10 Micropipette Beveler (Sutter Instrument, Novato, CA). The shRNA was injected using a nanoinjector (Sutter Instrument, Novato, CA). After all the shRNA was injected into the round window, the pipette was left in place for 30 seconds before removal. The muscles and fat tissue were covered, and the skin was closed with a polypropylene suture. The mice were kept on the warm bedding heated by a heating pad for recovery before returning to their mother. Total surgery time did not exceed 15 min.

### Voltage-clamp recording of hair cell mechanoelectrical transducer (MET) current

Patch-clamp experiments were performed in the standard whole-cell mode using an Axopatch 200B amplifier (Axon Instruments). Patch electrodes were pulled with a horizontal puller (Sutter Ins. Navato, CA) and had a resistance of 2–3 MΩ when filled with pipette solution consisting of (in mM) 135 CsCl, 10 HEPES, 2.5 EGTA, 0.25 CaCl_2_, MgCl_2_, 4 MgATP, and 0.4 Na_2_GTP (pH adjusted to 7.3 with CsOH). The bath solution consisted of (in mM) 130 NaCl, 3 KCl, 1 MgCl_2_, 10 HEPES, 2.5 CaCl_2_, and 10 glucose (pH was adjusted to 7.3 using NaOH). Currents were sampled at 20 kHz and filtered at 2 kHz. Voltages were not corrected for a liquid junction potential. No leak current subtraction was performed. Cells were held at −80 mV. All electrophysiological experiments were performed at RT (21-22°C). We used stepwise and sinewave mechanical stimulation of IHC bundles through a piezo-driven fluid-jet stimulator to record IHC mechanoelectrical transducer (MET) currents. We represented the bundle displacement in the form of applied piezo-driven voltage. Bundle displacement was not calibrated for each cell because of variations in stimulating probe positions relative to the stimulated hair bundle.

### Data analysis

The ABR and DPOAE data were analyzed using GraphPad Prism 7 (GraphPad Software, San Diego, CA, US) and OriginPro 2020 (OriginLab Corp., Northampton, Mass, US). Three-way ANOVA was used to analyze the ABR and DPOAE threshold. Two-way ANOVA was applied for the ABR wave 1 amplitude/latency data, synapse count, and HCs count. Significance was assumed at a *p*-value of 0.05 in all statistical analyses.

## Acknowledgments

We thank members of our laboratory for their comments on this manuscript. This work was supported by grants from the National Institutes of Health (DC016099, DC05135, AG051443, DC015252, and AG060504) to E.N.Y.

## Author Contributions

E.N.Y and BF designed the research, Y.C. M.C.P.F., J.H.L, J.L, S.P, M.K, B.P, J.K, J.C, L.L, MAG, and E.N.Y., analyzed data and wrote the manuscript. All authors read and approved the final manuscript.

## Author Information

Correspondence should be addressed to E.N.Y., the Program in Communication Science, the Department of Translational Neuroscience, the College of Medicine, Phoenix AZ, University of Arizona Phoenix, AZ.

---

**Supplement S1.**
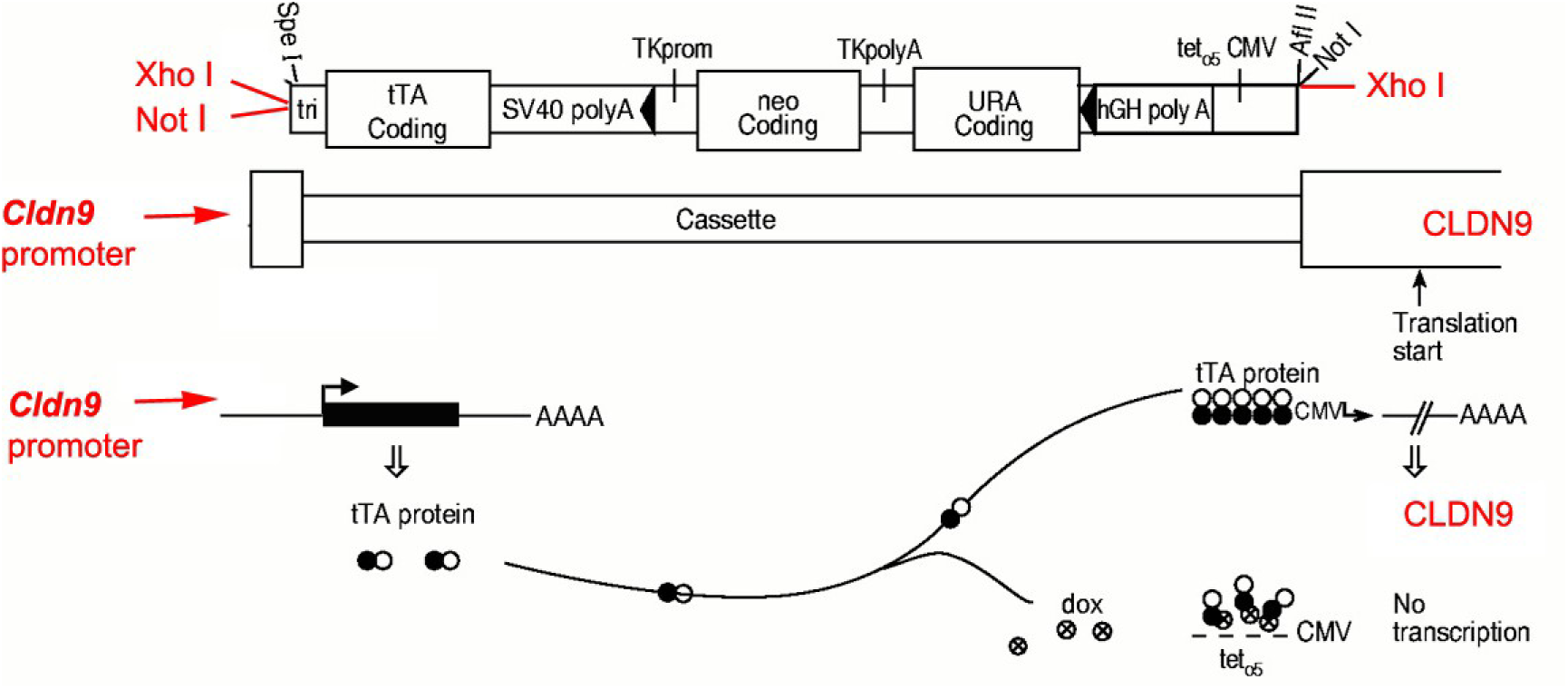
Schematic diagram of the construct used to regulate translation of CLDN9. We will use the tTA (“tet-off”) cassette developed by Seeburg and Adelman for expressing the SK3 channel {Bond, 2000 #182} (Bond et al., 2000) and modified to include unique *Not* I and *Xho* I cloning sites at each end of the cassette. The cassette contains three motifs: i.) tTA preceded by the tri-partite leader sequence that confers highly efficient translational initiation followed by the SV40 polyA and transcriptional termination sequence, ii.) bacterial neomycin (G418) resistance gene driven by the herpes simplex virus thymidine kinase (TK) promoter and the yeast URA3 gene; these selectable markers are flanked by loxP sites and followed by the human growth hormone (hGH) polyA and transcriptional termination sequence, iii.) six copies of the tet operator (tetO6) fused to the minimal cytomegalovirus (CMV) promoter. The native *Cldn9* promoter drives the expression of tTA, which in CLDN9 from the tetO6/CMV promoter. In the presence of Dox (crossed circles), tTA is bound and cannot bind to the tetO6/CMV promoter, thereby turning off the transcription of CLDN9.

**S2.**
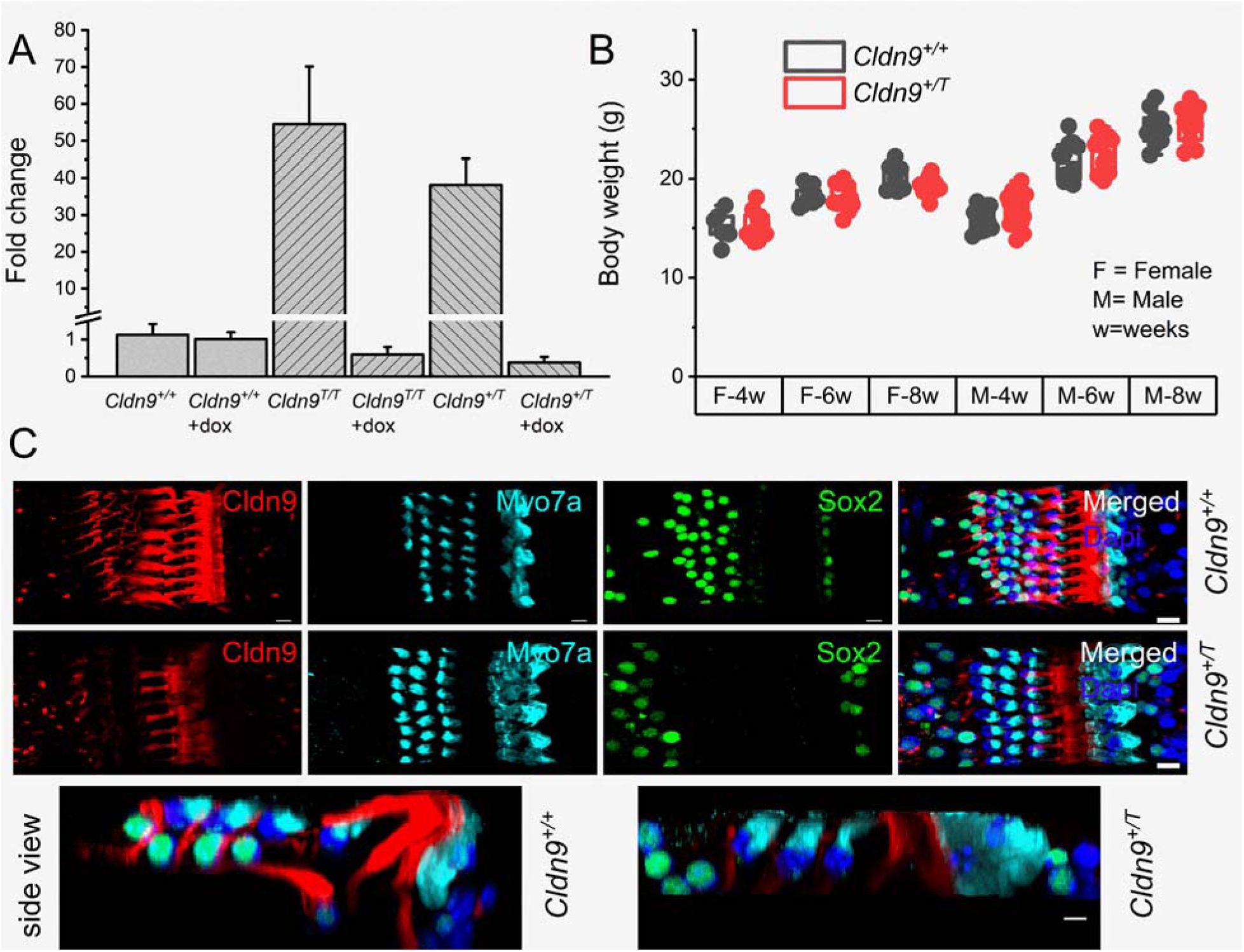
**A.** Quantitative RT-PCR of *Cldn9* transcripts from cochlear tissue from six groups of animals, including *Cldn9^T/T^*, *Cldn9^+/T^* with and without dox treatment compared with WT littermates with and without dox treatment. **B.** Body weight measurements of female and male mice from dox-treated *Cldn9^+/T^* and *Cldn9^+/+^* groups were recorded at 4, 6, and 8 wks old. **C.** The immunostaining of *Cldn9* in 8-wk old WT (*Cldn9^+/+^*) mouse cochlea*Cldn9* (red), IHC stained myosin7a (cyan) and supporting cells stained Sox2 (green) and Dapi (blue) for nuclear stain. Scale = 10 *μ*m.

**S3.**
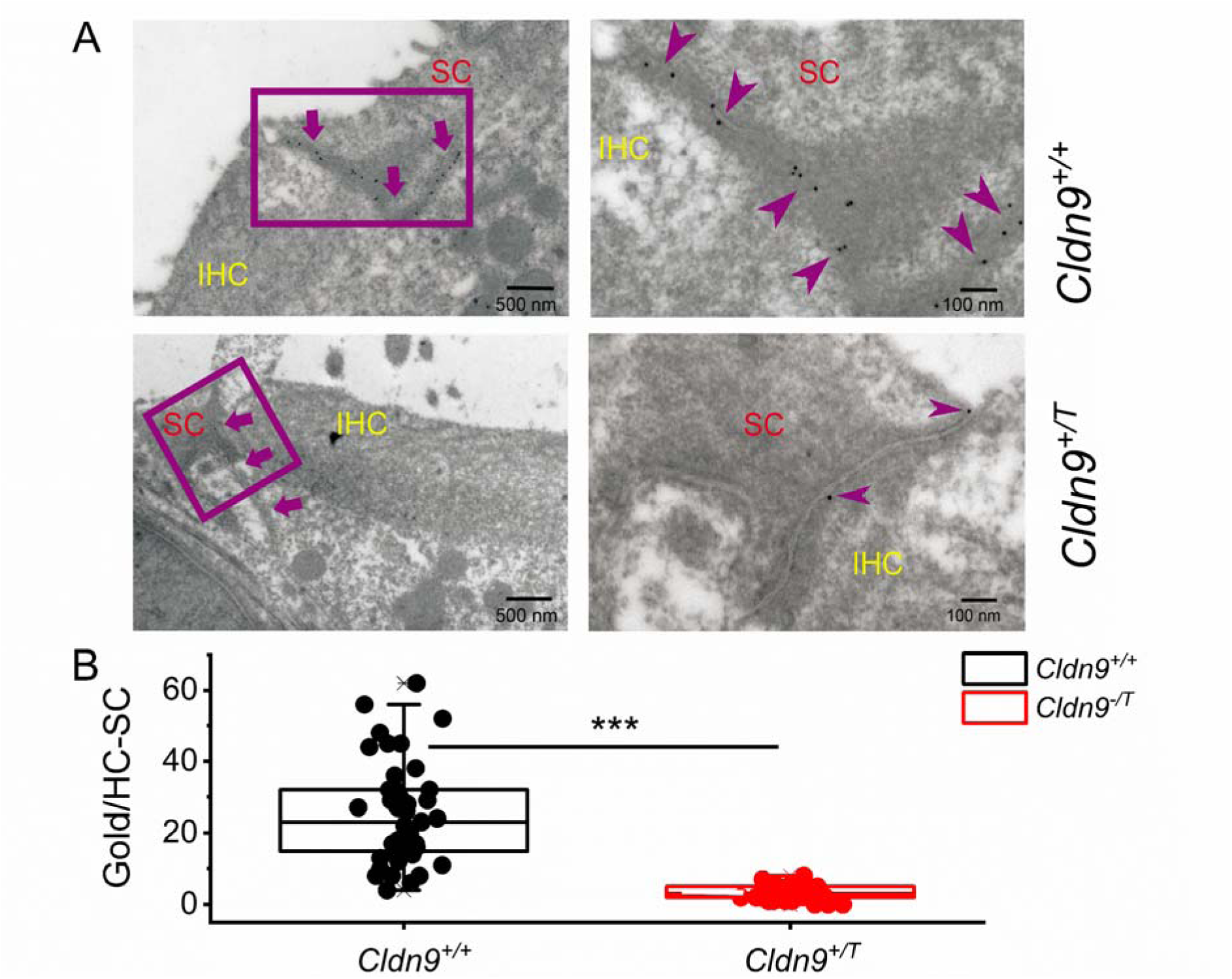
Immunogold localization of Cldn9 in *Cldn9^+/T^* and *Cldn9^+/+^* mice. **A.** Cldn9 expression sites between inner hair cells (IHCs) and supporting cells (SCs) were examined with immunogold electron microscopy with the post-embedding technique. Secondary antibodies are conjugated to 16-nm colloidal gold particles (arrows). Gold particles were noted at the junctions between SC and IHCs in *Cldn9^+/+^* mice cochlea (indicated). *Cldn9^+/T^* mice had reduced gold particles. **B.** Summary of gold particle counts between IHC and SC between Cldn9^+/+^ and Cldn9^+/T^. Mean gold particles (mean±SD) for *Cldn9^+/+^*= 25±14 (n = 41 from 3 cochleae) and for *Cldn9^+/T^* = 3±2 (n = 41, from 4 cochleae), *p = 2.8X10^-15^*.

**S4.**
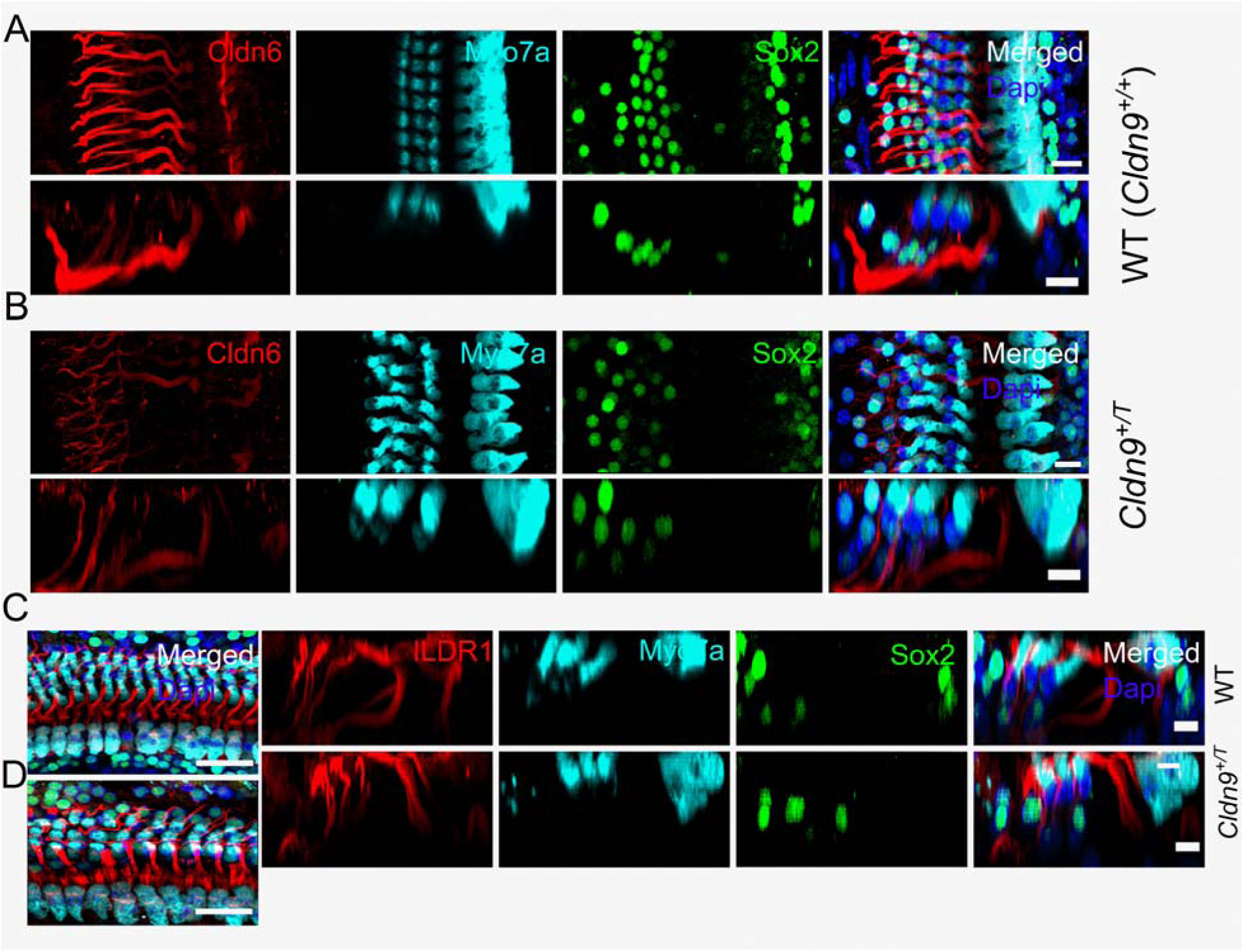
The expression of cldn6 and ILDR1 in the organ of Corti. **A-B,** The immunostaining of Cldn6 (red) in the mouse cochlea from *Cldn9^+/+^* (wildtype, WT) and C*ldn9^+/T^*. The lower Panel is the side view of the cochlear section. HCs were labeled with Myo7a (cyan) supporting cells with Sox2 (green) and Dapi-stained (blue) nuclei. The levels of Cldn6 were reduced in the *Cldn9^+/T^* cochlea. **C-D**, The immunostaining of ILDR1 (red) in the mouse cochlea from *Cldn9^+/+^* and C*ldn9^+/T^*. The lower Panel is the side view of the cochlear section. HCs were labeled with Myo7a (cyan) supporting cells with Sox2 (green) and Dapi-stained (blue) nuclei. The levels of ILDR1 were increased in the *Cldn9^+/T^* cochlea. Scale bar = 10 *μ*m.

**S5.**
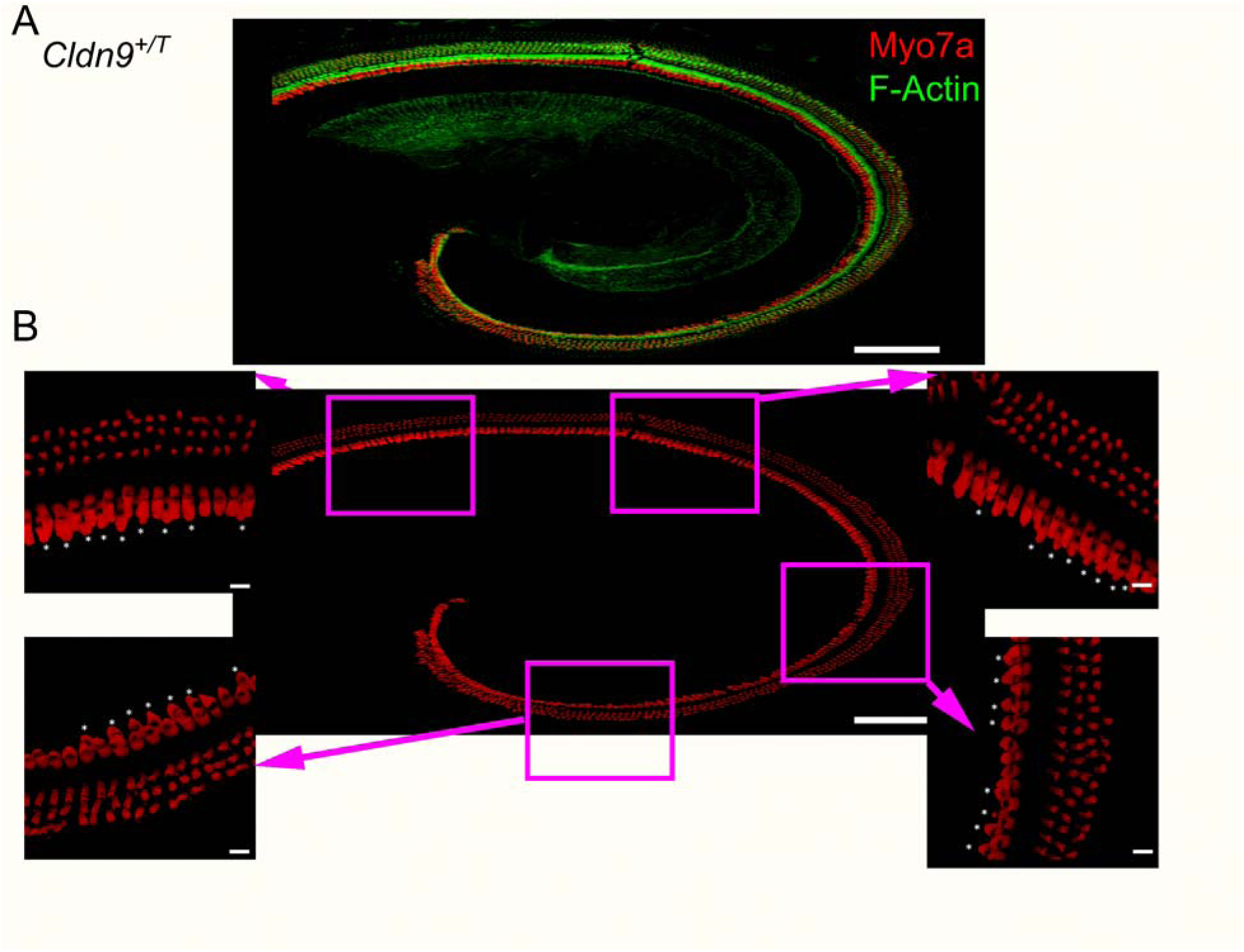
The ectopic HCs can be seen along the cochlear apicobasal contour. **A,** The immunostaining of Myosin 7a (red) and F-actin (green) in C*ldn9^+/T^* mouse cochlear apical segment. **B**, Panel shows smaller segments magnified to clarify and detect multiple ectopic HCs. HCs were labeled with Myo7a (red). Scale bar = 10 *μ*m.

**S6.**
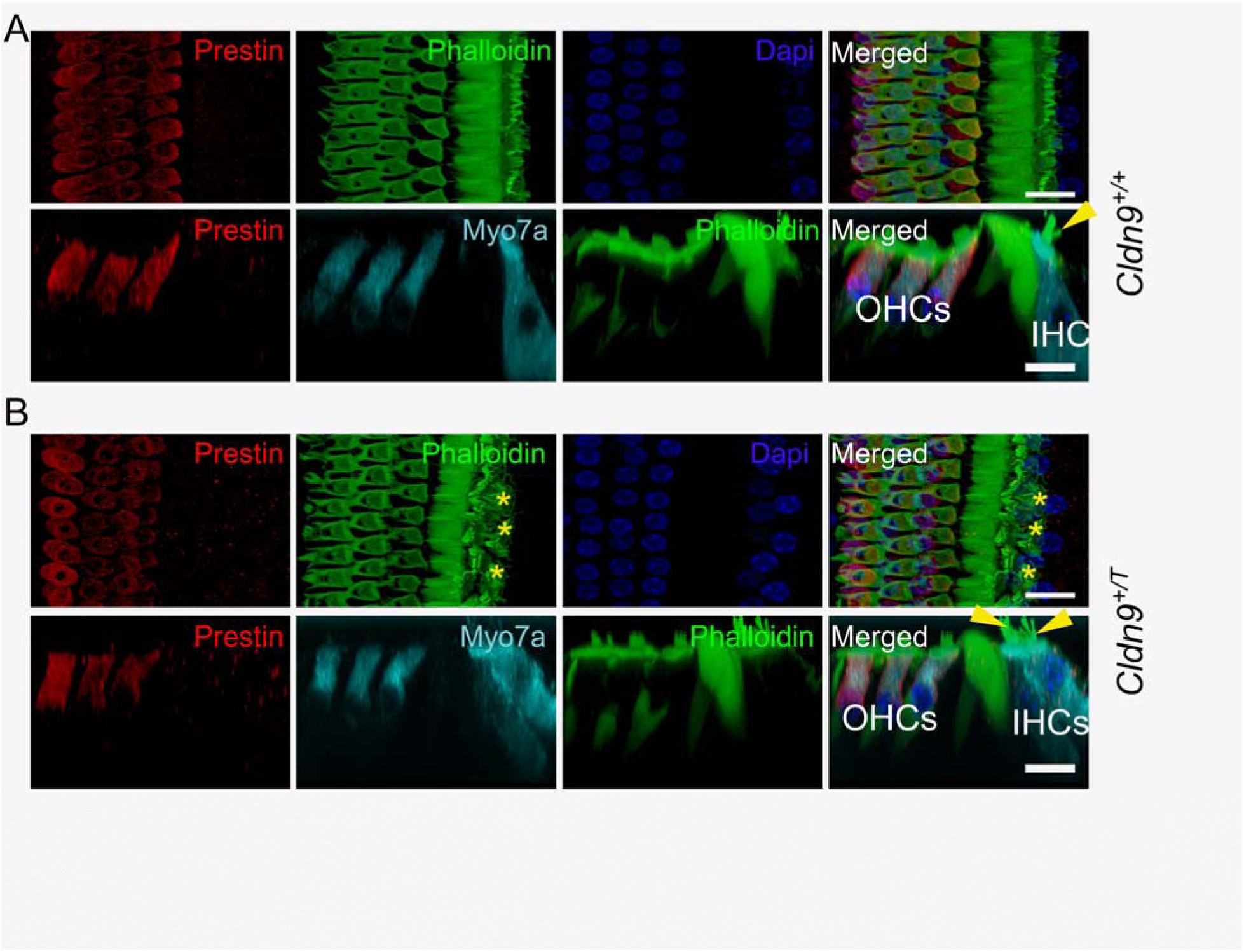
The ectopic HCs stain positively to Myosin 7a but negatively to prestin antibodies. **A-B,** The immunostaining of prestin (red) in the mouse cochlea from *Cldn9^+/+^* (wildtype, WT) and C*ldn9^+/T^*. The lower Panel is the side view of the cochlear section. HCs were labeled with Myo7a (cyan) phalloidin-stained actin (green) and Dapi-stained (blue) nuclei. Scale bar = 10 *μ*m.

**S7.**
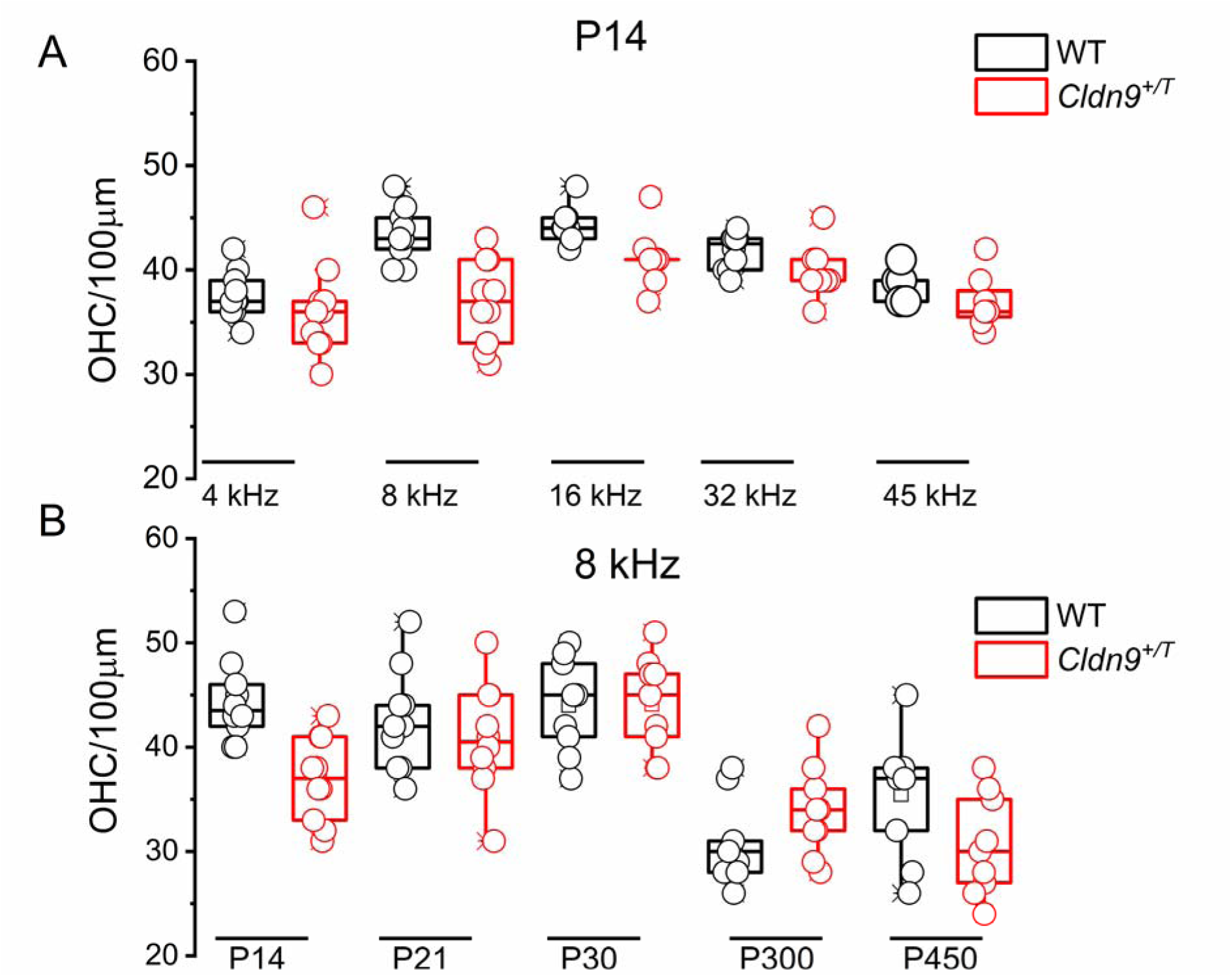
**A.** Summary data of Quantification of OHCs count at ∼4, 8, 16, 32, and 45-kHz placed cochlear map at postnatal days (P14) in *Cldn9^+/+^*(WT, in black symbols) compared with *Cldn9^+/T^* (in red symbols) mice. Each data point is the mean of 4-blind counts. **B.** At 8 kHz cochlear placed-map, OHC counts were changed for P14, 21, 30, 300, and 450 mice. OHC numbers remained relatively constant (P14-30) until P300-450) when they were reduced.

**S8.**
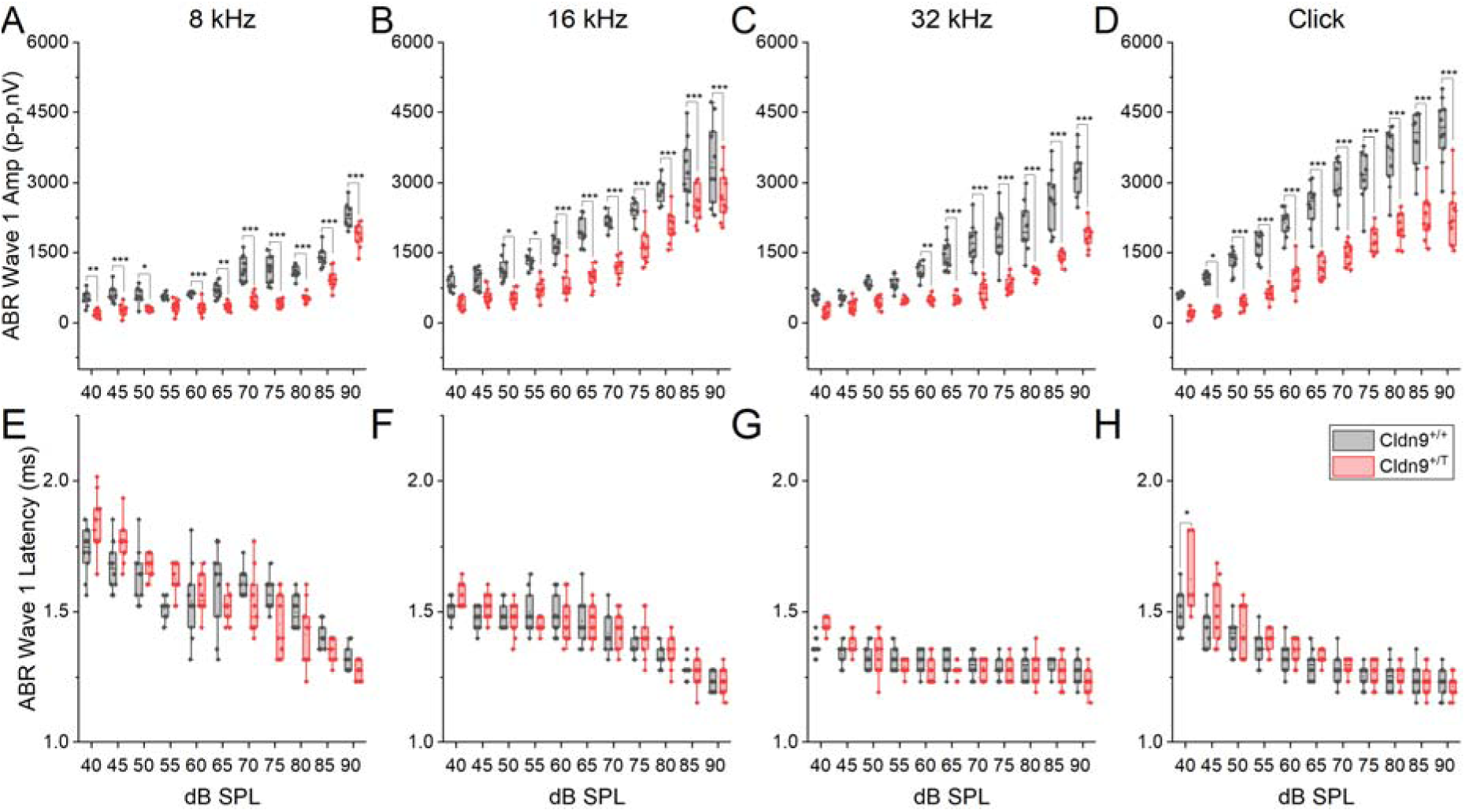
The ABR wave one amplitude and latency in *Cldn9^+/^*^T^ and *Cldn9^+/^*^+^ mice. **A-D.** ABR wave 1 amplitude was decreased, while the latency varied in *Cldn9^+/^*^T^ mice compared to *Cldn9^+/^*^+^ mice when measured at 8 kHz, 16 kHz, 32 kHz, and click **(E-H).** * P < 0.05, ** P < 0.01, *** P < 0.001.

**S9.**
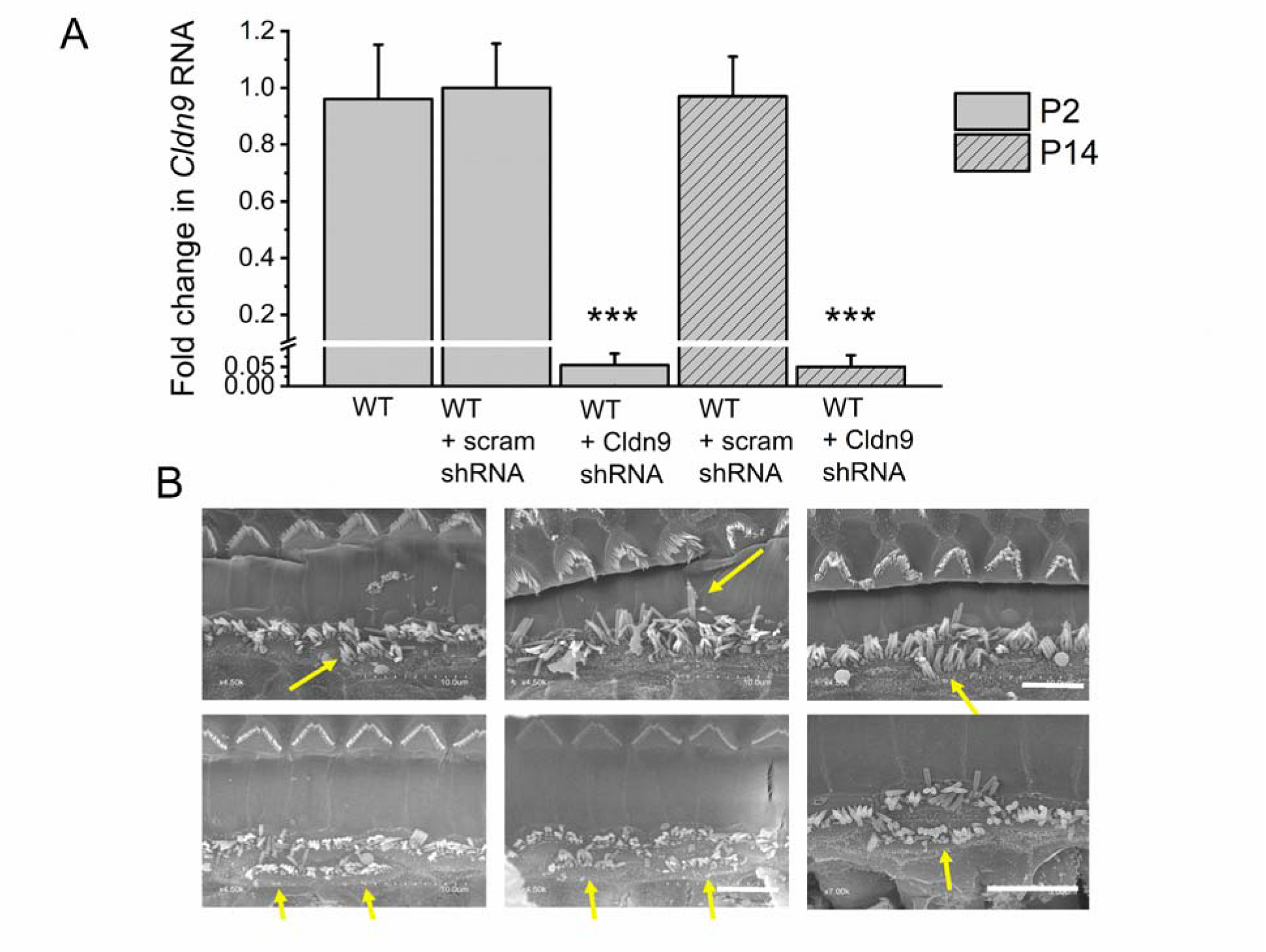
**A. Quantitative RT-PCR of *Cldn9*** transcripts from cochlear tissue from WT un-injected, scrambled (scram), and shRNA P2- and P14-old-injected mice. **B.** Examples of SEM of the shRNA-injected cochlea (age of samples at the harvesting time (P24) for imaging and the age at shCldn9 injection P2). Yellow arrows show ectopic IHC bundles.

**S10.**
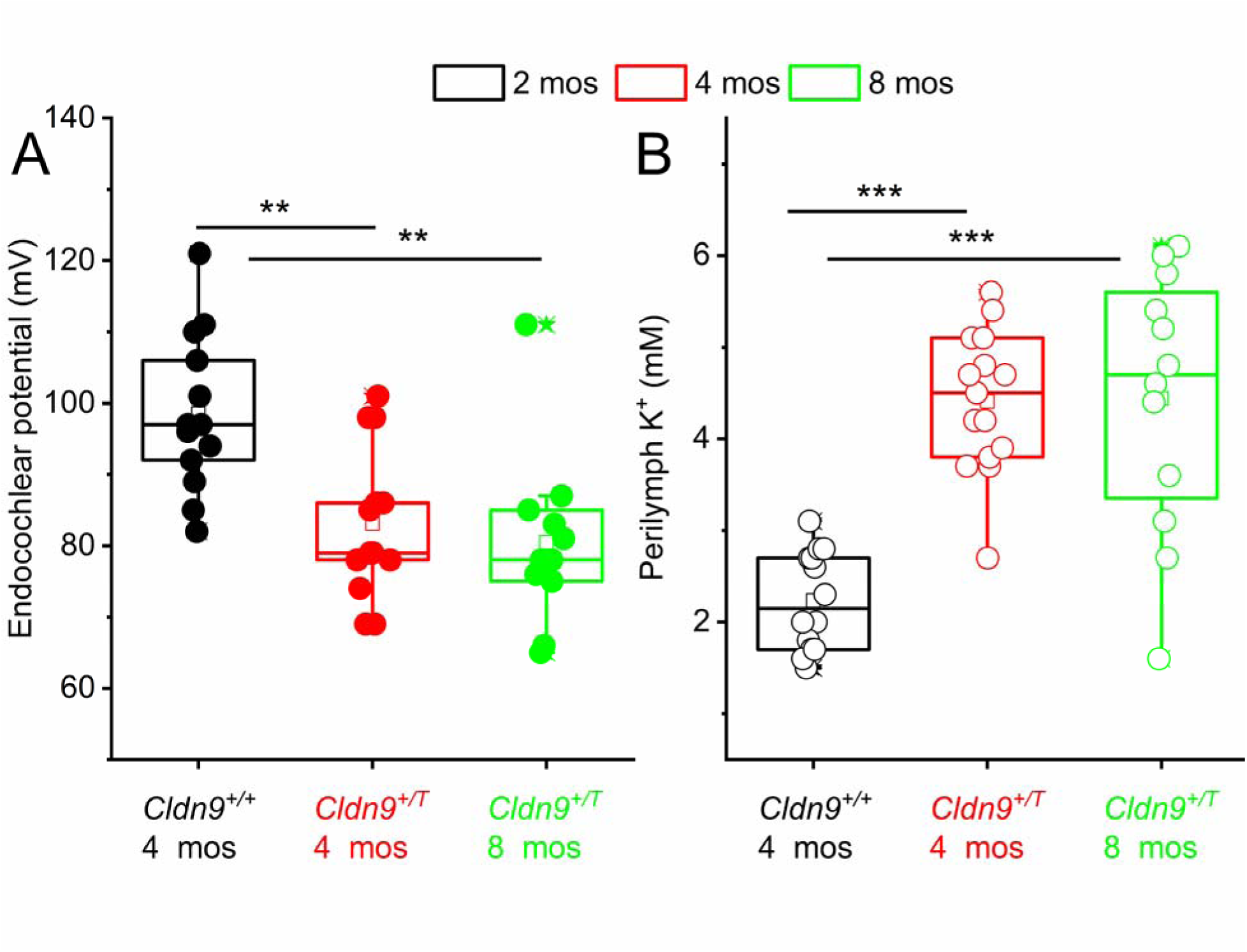
The endocochlear potential (EP) and perilymph K^+^ in *Cldn9^+/+^* and *Cldn9^+/T^*mice. **A.** Measurement of the EP at the basal turn of the cochlea, in 4-month-old *Cldn9^+/+^* = 99±11 mV (n=13), *Cldn9^+/T^* = 83±11 mV (n=13) and at 8-month-old for *Cldn9^+/T^ =* 80±12 (mV) (n=11). There were significant differences at *p*<0.05 level for comparison between *Cldn9^+/+^* vs. *Cldn9^+/T^*F(2,34)=(9.3) *p* =5.9*X10^-4^*. *Post hoc* comparisons using the Tukey HSD test indicate *Cldn9^+/+^* vs. *Cldn9^+/T^* at 4 months (*p*=4.0X10^-3^); *Cldn9^+/+^* (4 months) vs. *Cldn9^+/T^* (8 months) (*p*=1.2X10^-3^); are significantly different. **B.** Recordings of perilymph K^+^, in 4-month-old *Cldn9^+/+^* = 2.2±0.5 mM (n=14), *Cldn9^+/T^* = 4.4±0.8 mM (n=15) and at 8-month-old for *Cldn9^+/T^=* 4.4±1.4 (mM) (n=12). There were significant differences at *p*<0.05 level for comparison between *Cldn9^+/+^* vs. *Cldn9^+/T^* F (2,36)=(24.4) *p* =1.6*X10^-7^*. *Post hoc* comparisons using the Tukey HSD test indicate *Cldn9^+/+^* vs. *Cldn9^+/T^* at 4 months (*p*=1.1X10^-6^); *Cldn9^+/+^* (4 months) vs. *Cldn9^+/T^* (8 months) (*p*=2.3X10^-6^); are significantly different.

